# Robust parameter estimation and dose-fractionation for ptychographic single-particle analysis at near-atomic resolution

**DOI:** 10.64898/2026.11.20.768111

**Authors:** Tizian Lorenzen, Daniel Mann, Jasenko Zivanov, Max Leo Leidl, Andreas Jehle, Vishal Kumar, Benedikt Diederichs, Franziska von Hammerstein, Mikhail Shein, Anne K. Schütz, Henning Stahlberg, Carsten Sachse, Knut Müller-Caspary

## Abstract

We present a complete workflow for the determination of biomolecular structures by single-particle cryogenic electron microscopy using ptychography. To account for the prominent dose damage in biological specimens dose-fractionated data acquisition and processing are introduced. Automated preprocessing techniques estimate the probe shape and scanning geometry before the ptychographic reconstruction by utilising the shadow image inside the Ronchigram. Standard ptychographic single-particle analysis is performed using tobacco mosaic virus, apoferritin and hepatitis B virus capsids. We finally give a demonstration of dose-fractionated ptychography utilising multiple scans of the same area at low dose reaching near-atomic resolution (3.3 Å), allowing for the automated building of atomic models.

## 1 Introduction

With electron optics achieving resolutions in the sub-Å range, transmisssion electron microscopy (TEM) is well established as a central technique to solve structures down to the atomic level in both physical and life sciences [1–3]. In recent years, these research fields experienced a dramatic evolution owing to innovations in detector hardware [4] and image processing [5–7]. In the physical sciences, ultrafast diffraction cameras [8–10] recording at frame rates of a few to more than one hundred kHz paved the way to efficient phase contrast imaging in scanning TEM (STEM). In particular, recording a full diffraction pattern at each position of a scanning probe provides a four-dimensional dataset that, by combining simultaneously acquired real and diffraction space information, allows for advanced phase retrieval via ptychographic algorithms [11]. Besides largely eliminating the effect of the electron optics [12], and thereby achieving superresolution down to the limits of thermal vibrations of atoms [13], electron ptychography employing 4D-STEM data is a highly flexible and versatile computational imaging technique with excellent contrast transfer and dose efficiency [14–16].

Recently, STEM employing focused probes scanning over the specimen (as shown schematically in Fig. 1 a) has found pilot applications in cryogenic electron microscopy (cryo-EM), in contrast to conventional TEM (CTEM) using parallel illumination [17–20]. In this respect, phase-contrast 4D-STEM methods gained attraction due to their monotonous contrast transfer, and the degree of freedom to employ scattering models of different complexity during the reconstruction. These models range from weak phase objects [21], as used in cryo-TEM, over phase and arbitrary, complex objects accounting for single scattering [5], to objects taking interactions with thick specimens into account, namely within multislice and scattering matrix approaches [22–24]. Regarding applications of phase-contrast STEM-imaging in structural biology, near-atomic resolution on single-particle specimens was demonstrated using integrated differential phase contrast (iDPC) imaging employing four-segment detectors [18]. In addition, spatial resolutions in the range of 5.8 and 8.4 Å [25] have been achieved using electron ptychography employing pixelated detectors. The rapid evolution of cryo-electron ptychography [17, 25, 26] was mostly based on the direct transfer of algorithms from its physical science counterpart rather than accounting for the inherent low signal-to-noise conditions in low-dose cryo-EM imaging. It can be anticipated that further progress in this field, such as achieving a spatial resolution sufficient for molecular model building, requires the development of a workflow dedicated to overcoming the specific challenges of imaging ice-embedded biological specimens.

**Fig. 1.**
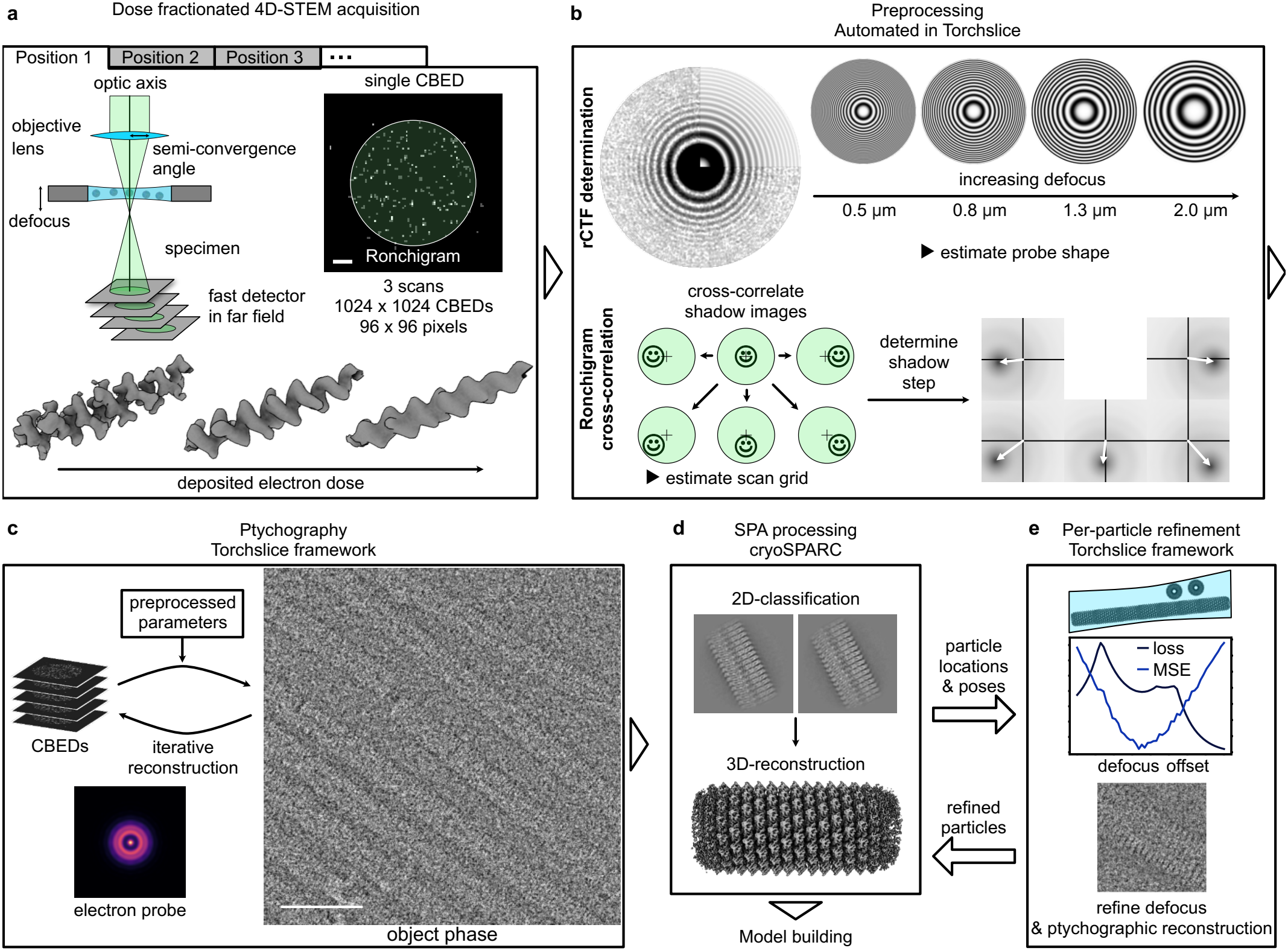
Full 4D-STEM single-particle analysis workflow. (a) Schematic representation of the experimental setup and an exemplary convergent beam electron diffraction pattern with the highlighted Ronchigram (length of scalebar: 1 mrad). At each specimen location multiple scans are acquired, thereby obtaining dose fractions to compensate for the effect of dose damage (schematic representation). (b) During preprocessing the probe shape and scan grid are estimated (c) which is used to initialise the ptychographic reconstruction (length of scalebar: 50 nm). (d) Utilising these reconstructions, single-particle analysis is performed to determine 3D maps which can be used for model building. (e) The obtained 3D-maps can then be employed for per-particle defocus refinement.

Here two main avenues appear. Firstly an automated processing pipeline needs to be developed eliminating the need for manual, and thus subjective, parameter tuning. Currently, these manual steps demand expert skills and limit the application of cryo-ptychography to a small user group in the biological sciences. Secondly, more advanced processing steps are to be developed in order to deal with the low signal-to-noise data, the pronounced irradiation-induced damage [27] and specimen movement [28]. In that respect, developing 4D-STEM dose-fractionation strategies is crucial to improve spatial resolution [29], in addition to the need for the highly accurate determination of the probe wave function, the independent retrieval of the scan raster geometry under ultra-low-dose conditions, as well as developing refinement concepts of the 3D density, e.g., via per-particle focusing [30].

This work introduces a comprehensive ptychographic workflow covering all stages from dose-fractionated 4D-STEM data acquisition at accumulated doses of 25 e^*−*^ / Å^2^ to the ptychographic retrieval of 3D densities. Standard ptychographic SPA is demonstrated using tobacco mosaic virus (TMV), apoferritin (ApoF) and hepatitis B virus capsids (HBV). The dose-fractionated ptychographic image processing workflow was used to determine the cryo-EM structure of TMV at a spatial resolution of 3.3 Å. We finally subject the obtained densities to model-building in order to obtain the molecular structure of proteins [31].

## 2 Results

### 2.1 Overview of ptychographic SPA workflow

The full workflow of cryo-ptychography for single-particle analysis (SPA) is shown exemplarily for a vitrified tobacco mosaic virus (TMV) specimen in Fig. 1 and consists of five major steps - data acquisition, pre-processing, ptychographic reconstruction, single-particle analysis and the optional per-particle defocus refinement. It starts with the acquisition of dose-fractionated 4D-STEM data by sequential recording of cartesian scans of 3*×*8 e^*−*^ / Å^2^ fluence according to Fig. 1 a.

As a prerequisite for stable and reliable specimen retrieval at low-dose settings, the procedure developed in Fig. 1 b serves to prepare the ptychographic reconstruction with an initialisation of crucial microscope-related parameters as accurately as possible. To this end, the central parts of the diffraction patterns, the Ronchigrams, are evaluated in real and Fourier space, respectively. As Ronchigrams recorded out of focus contain a shadow image projection of the illuminated specimen region at a location characteristic for the position of the scanning probe, the correlation of neighbouring Ronchigrams is employed to obtain the pitch, shear and rotation of the scan raster as shown schematically in the lower part of Fig. 1 b. The ability to estimate the scan parameters from the data itself is especially important in cryo-EM ptychography since standard methods for scan step calibration (e.g. gold or silicon lattice planes) are performed at higher magnifications where lattice planes can be resolved (typically using scan steps smaller than one Ångstrom) and then extrapolated to lower magnifications. Furthermore, standard methods for defocus estimation of 4D-STEM data using e.g. parallax imaging [19, 32] or even ptychographic focal series (Suppl. Fig. 2) produce wrong defoci when the scan step is not accurately known.

By further analysis of the Ronchigram’s contrast transfer function (rCTF) using the Fourier transformed Ronchigrams, the actual focus setting and residual astigmatism are estimated (Fig. 1 b top). During preprocessing, the bright field movement arising from residual scan position-dependent beam tilts is additionally mapped (Supplementary Note 1) and scan positions on thicker supports are discarded.

After preprocessing and initial defocus estimation, GPU-opitimized ptychographic reconstructions are performed as shown in Fig. 1 c, while employing a patch-wise iterative update scheme and efficient hyper-parameter tuning. Throughout this work, we use the term *reconstruction* for the ptychographic result, and *3D-reconstruction* for the generation of 3D densities. With the set of individual phase reconstructions, the established single-particle workflow, as available in e.g. cryoSPARC [7], is used to obtain a 3D density (Fig. 1 d). Since particle position and orientation have been estimated, further reference-based refinement and perparticle defocus optimisation can be used to compute improved particle images, which can be can be fed back to cryoSPARC to enhance the resolution of the 3D map (Fig. 1 e).

In the following sections, these steps are further detailed with the aim to enable straightforward implementation of the full procedure in contemporary cryo-TEM laboratories equipped with 4D-STEM acquisition hardware. For further theoretical details and mathematical expressions the reader is referred to the *Methods* section.

### 2.2 Defocus and scanning geometry estimation using the Ronchigram

For the estimation of the scanning geometry and the defocus the shadow image appearing in Ronchigrams recorded with a strongly defocused probe as illustrated in Fig. 2 a,b is utilised. Because the shadow images in Fig. 2 b resemble a bright field CTEM image of the illuminated region in Fig. 2 a with a magnification proportional to the reciprocal of the defocus [33], the (local) defocus can be estimated from the contrast transfer function of this shadow image and the scanning geometry from its shift during scanning.

**Fig. 2.**
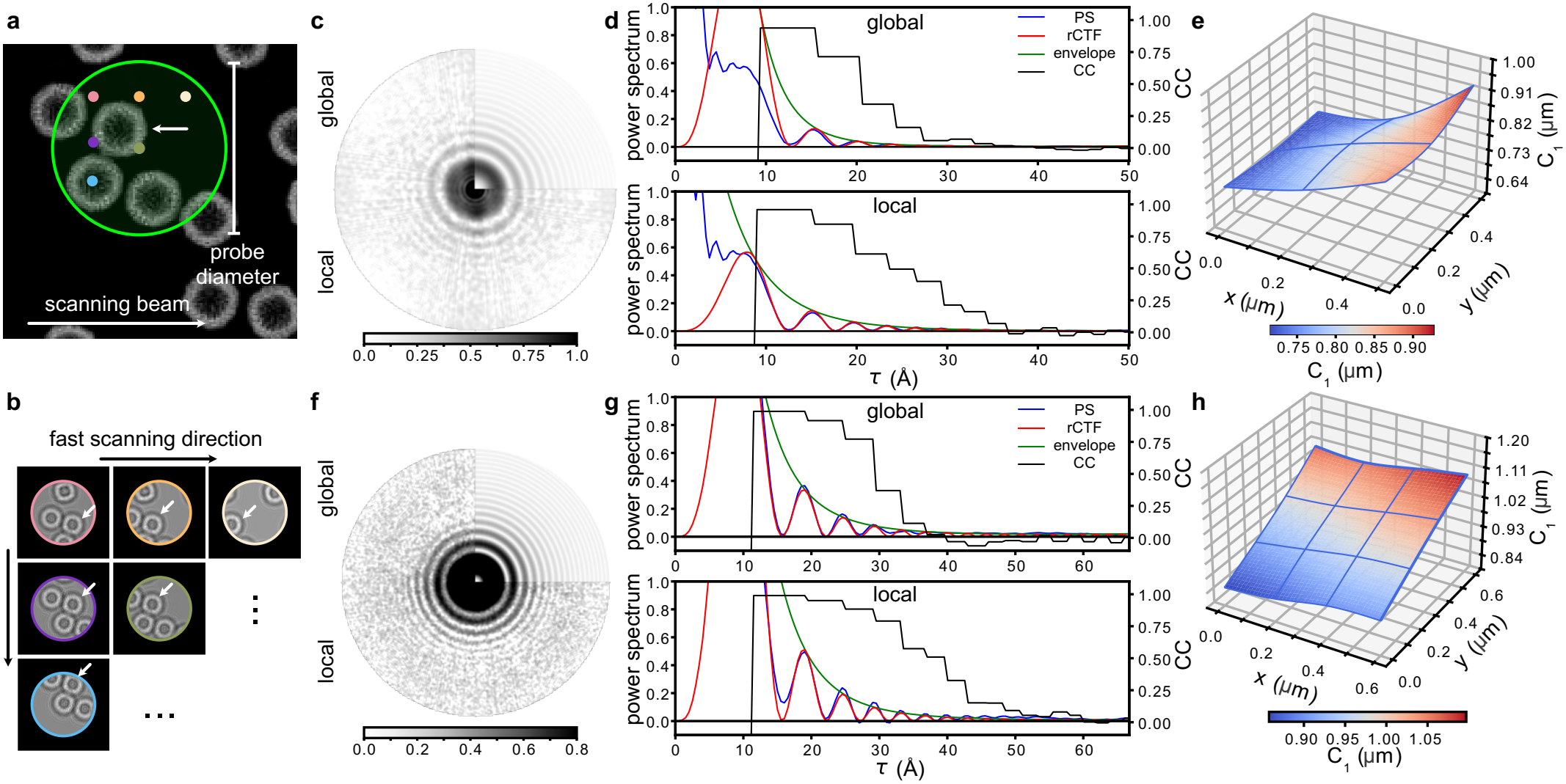
Local defocus estimation using the Ronchigram contrast transfer function (rCTF). (a) A defocused electron probe produces a shadow image of the specimen in the (b) diffraction pattern. The green circle indicates the area illuminated by a single probe position (simulated data). This shadow image is affected by the rCTF as visible by the fringes and moves with the beam, the arrows marking the same particle in the specimen and the shadow images. (c) Background-subtracted mean power spectrum of the shadow images without (global) and with re-scaling accounting for the local defocus variation (square root scale) obtained from a scan of an apoferritin sample. The fitted rCTF is displayed in the top right corner. (d) Diagnostics plot displaying the equiphase average of the same power spectrum (blue), the rCTF (red), its envelope (green) and the correlation coefficient (CC, black). (e) Local defocus landscape of the imaged area. Crossings of the blue lines indicate the position of the 3*×*3 grid of control points. The power spectra were calculated in patches of 64*×*64 scan positions with an overlap of 50%. The analysis is repeated for a scan of a standard cross-grating sample employing a Timepix4 detector in f-h.

To measure the defocus of the STEM probe, we evaluate the power spectra of the shadow images in the Ronchigrams. For weak scatterers as in the present work on SPA, the periodicity of the rCTF-pattern in radial direction and potential anisotropies are characterisitic for defocus and astigmatism as shown in Fig. 1 b which has been described quantitatively [34]. Note that only the power spectra of Ronchigrams are considered. This means that the phase in Fourier space is disregarded, such that the focus measurement is agnostic to the movement of the shadow images during scanning and hence fully independent of the knowledge about the scan pattern. As a consequence, the method is not restricted to cartesian scan patterns but can be applied to arbitrary scans or even single diffraction patterns.

In our rCTF determination approach, the power spectra are averaged within patches of scan positions, e.g. 64 x 64 at 1 µm defocus as in Fig.2, and the defocus is modelled using cubic B-splines [35]. After local defocus determination the power spectra are rescaled to account for these local defocus variations [35, 36]. Importantly, the iterative optimisation procedure does not require recalculation of the power spectra. As a consequence, the number of convergent-beam electron diffraction patterns (CBEDs) only contributes to the computational expense of the calculation of the power spectra whereas the computational demand for the rCTF determination only depends on the detector size and number of patches.

This is demonstrated in Fig. 2 c-e using a scan of a tilted apoferritin sample. The comparison of the background-subtracted power spectrum before (Fig. 2 c, top left) and after the rescaling procedure (Fig. 2 c, bottom) reveals that additional rings are visible indicating the improved determination of the local defocus. The same pattern is correctly reproduced by the modeled rCTF shown in the upper-right quadrant. Furthermore, the profiles in Fig. 2 d show an excellent agreement between the rCTF model (red) and the experimental background-subtracted power spectrum (PS, blue) in terms of azimuthal averages. The azimuthal averaging is performed along isolines of the rCTF to remove the blurring effect of astigmatism, referred to as equiphase averaging [37]. Significant correlation coefficients (black) extend to much higher frequencies τ in the case of local defocus estimation than for the global one. Using a threshold of 0.25, the limit up to which rings can be detected increases by 25 % from τ = 24 Å (global) to τ = 30 Å (local). As expected from the experimental setting, the overall tilt of the foil is seen in the map in Fig. 2 e and measured to 17.4^*°*^, which is in good agreement with the introduced specimen-stage tilt of 20^*°*^.

Because the method is based on the averaged power spectra of the shadow images it greatly profits from high sampling of the Ronchigram by the employed diffraction camera. While the primary detector used in this study is a Dectris Arina high-speed camera with 96 x 96 pixels, recent Timepix4 detector technology [38, 39] comprising 512*×* 448 pixels is becoming available, whose event-based detection scheme furthermore relaxes constraints on the scan speed for frame-based cameras. We thus additionally performed an analogous evaluation on a 4D-STEM scan of a tilted gold on amorphous carbon specimen recorded on a Timepix4 chip. Analysis of the power spectrum in Fig. 2 f shows an increased number of rings compared to Fig. 2 c. Note that the improvement when evaluating local scan patches is stronger than in the prior example, which is a result of the increased semi-convergence angle (SCA, 9 instead of 6 mrad) and highlights the importance of local defocus estimation especially when using highly convergent probes. The improved recovery is particularly visible in the azimuthal averages in Fig. 2 g.

Having estimated the probe shape from the power spectra of the diffraction patterns, we now evaluate the movement of the shadow image during scanning to estimate the scanning geometry. This movement is revealed by tracking features across adjacent shadow images, e.g. by apoferritin particles (indicated white arrow in Fig. 2 b). Since the observed shift is linear in the scan step, the cross-correlations of neighbouring shadow images can be utilised to accurately estimate the scan raster geometry. Importantly, the low SNR in biological applications usually precludes the evaluation for each scan point individually [40]. As shown in Fig. 3 a, the reliability of estimation improves significantly by averaging tens to several thousands of cross-correlations of Ronchigrams from scan positions with the same relative probe positions and a clear cross-correlation peak becomes visible. Here, the distance of this peak from the centre contains information about the relation of the scanning step to the magnification of the shadow image while its azimuthal orientation yields the relative orientation of the scan and camera coordinate systems.

**Fig. 3.**
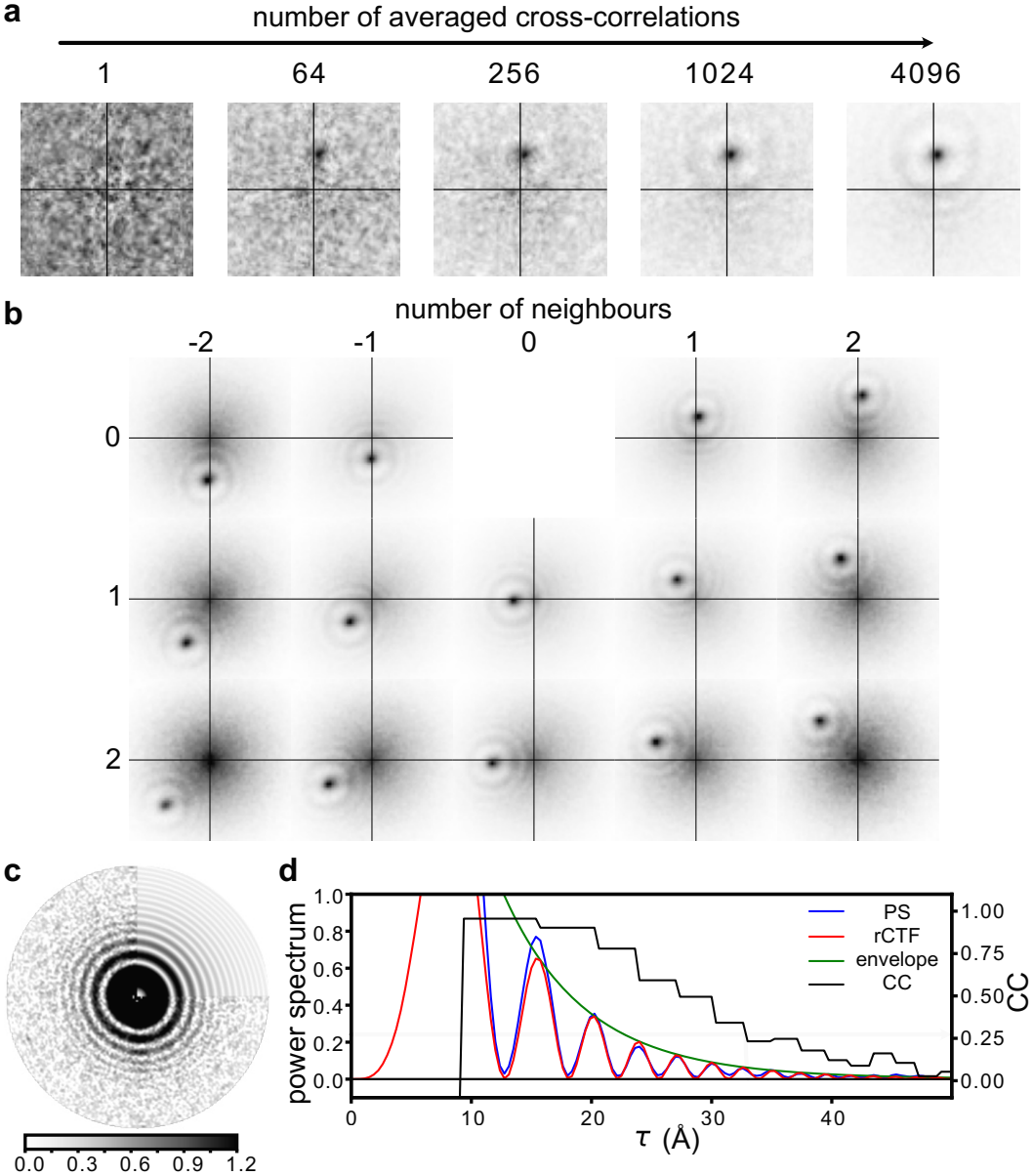
Scanning geometry estimation using the cross-correlation of neighbouring Ronchigrams. (a) Cross-correlation of CBEDs from neighbouring probe positions reveals the shifting in the shadow image. The SNR of these cross-correlations increases when averaging multiple diffraction patterns. Cross-correlation plots for a non-tilted grid with the numbers on the top and side indicate the neighbor order *n* in the fast and slow scan direction respectively. Shown are the central parts of the cross-correlations up to *n* = 2 in both directions. The black crosses indicate the center of the coordinate system. (c) Power spectrum (square root scale) of the shadow image recovered from the cross-correlation peaks and the fitted rCTF. (d) Radial average of the same area.

Note that nearest neighbours as well as higher-order neighbours should be included, as long as the two correlated Ronchigrams still exhibit common features. In practice, calculating the cross-correlations up to neighbour orders *n* = 2 or 3 in the fast and slow scan direction was sufficient, as shown in Fig. 3 b for a 4D-STEM scan across an untilted grid. The mean geometry of the scan raster in Fig. 3 b was determined to have a scan pitch of 19.6 and 19.8 Å in the fast and slow scan direction, respectively. Shear of the scan raster was found negligible here as the two scan axes incline an angle of 90.15^*°*^, furthermore an overall rotation of *−*0.42^*°*^ between the coordinate frames of the scan and the diffraction camera was determined. The ptychographic reconstruction shows the spherical apoferritin particles with high contrast (Suppl. Fig. 3).

In the presence of local defocus variations, the changes in shadow image magnification result in anisotropic smearing of the cross-correlation peaks. If such local defocus variations are present, the cross-correlations are calculated in scan patches and scaled to the same average magnification using the defoci determined using rCTF analysis, which allows the determination of the scanning geometry even in tilted specimen as shown in Suppl. Fig. 4. The impact on the cross-correlations is again significantly more pronounced in the dataset recorded at the higher SCA of 9 mrad (Suppl. Fig. 4 b).

Since the observed cross-correlation shifts are linear in both the scanning step and the focus-dependent magnification of the shadow image, the Ronchigram cross-correlations can as well be used to determine the defocus, astigmatism and the orientation of the camera when an accurate calibration of the scanning geometry is available.

A closer look at the cross-correlations in Fig. 3 b reveals a faint ring-shaped structure centred around the maximum, which deserves special attention. Under the assumption that two neighbouring Ronchigrams only vary in a shift, this modulation of the peak is given by the inverse Fourier transform of the power spectrum of the shadow images (see Methods). Importantly, this opens up an alternative method for retrieving the power spectrum of the shadow image and thus to gauge the defocus [33]. In particular, we calculate an average auto-correlation by shifting all cross-correlation peaks to the origin. The power spectrum is obtained by Fourier transformation and subjected to the rCTF determination procedure. As can be seen in Fig. 3 c the power spectrum determined in this way shows a reduced background and is well represented by the rCTF model. This excellent agreement is also visible in the azimuthal averages in Fig. 3 d where the reduced background results in a fit resolution of τ *≈* 33 Å.

### 2.3 Ptychographic reconstruction and SPA

The described estimates of the STEM probe and the scan raster are used to initialise the ptychographic reconstructions. In contrast to blind ptychography where both probe and specimen need to be retrieved from scratch, our approach does not only speed up convergence but also minimises the risk of getting stuck in local minima, given the noisy 4D-STEM data. Ptychographic reconstructions were performed using the automatic-differentiation based TorchSlice package [24, 41]. Note that rather than denoting a single approach, the term ptychography includes a broad range of methodologies which differ significantly in terms of the assumed interaction model between probe and specimen, whether direct or iterative schemes are used, which loss function is employed and whether physics-related parametrisations are used. While non-ptychographic probe and scan grid retrieval schemes introduced above would already allow for the deconvolution of probe and specimen *via* direct ptychography schemes [21, 42] reformulated to be independent from the scan grid [43], this study makes use of iterative concepts as they provide more flexibility as to probe refinement and noise models.

Briefly, we utilised a single scattering model comprising a complex object transfer function (OTF) and a single coherent probe, having been proven accurate for the single-particle specimens in simulation studies [44, 45]. Automated estimation of the learning rates is performed using hyperparameter optimisation on a small subset of the scans [46, 47]. As an important addition to the reconstruction, we include the shift of the bright field disk during scanning, which can be essential for datasets in which large fields of view are scanned (Supplementary Note 1). See Fig. 4 a-d for example reconstructions of TMV, apoferritin and HBV capsids discussed comprehensively below. After performing a fixed number of iterations the retrieved object phases are subjected to further SPA processing.

**Fig. 4.**
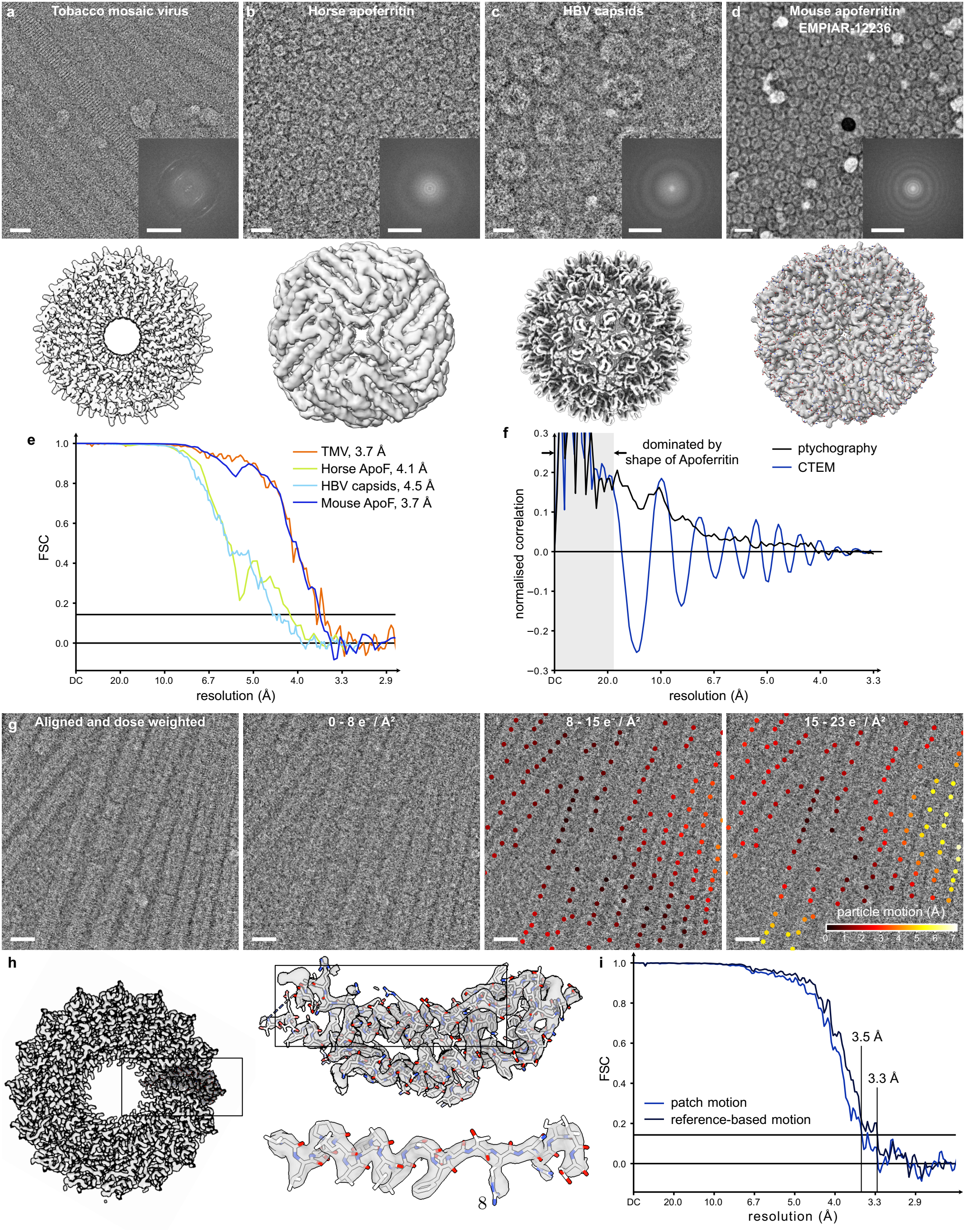
Standard and dose-fractionated ptychographic single-particle analysis. Ptychographic micrograph (top) and reconstructed 3D-map (botttom) using ptychographic single-particle analysis without dose-fractionation of (a) TMV, (b) horse spleen apoferritin, HBV capsids and (d) mouse apoferritin from EMPIAR-12236 (fitted model ID: PDB 8RQB) [25]. The insets show the power spectra of the micrographs. (e) FSC curves indicating resolutions of 3.7, 4.1, 4.7 and 3.7 Å (FSC=0.143 cutoff), respectively. (f) Normalised correlation between projected reference and particle patches for a ptychographic and a conventional TEM micrograph. (g) Producing a dose-fractionated micrograph by aligning and dose-weighting the reconstructions of the dose fractions visibly increases contrast. The absolute per-particle motion relative to the first fraction is shown for each particle. (h) Slab view of TMV with the built model after local resolution filtering (left), the asymmetric unit of TMV and a single alpha helix highlight the quality of the model (right). (i) FSC curves indicating a resolution of 3.5 Å using the motion-corrected micrographs and of 3.3 Å after reference based motion correction. Scalebars equal 20 nm and 1 nm^*−*1^.

The SPA processing is conducted using cryoSPARC, the exported ptychographic micrographs can be treated using standard workflows, e.g. by performing particle picking, 2D-classification and 3D-reconstruction. Importantly, however, no CTF determination and correction needs to be included as the utilised object phase is directly proportional to the projected potential of the specimen.

### 2.4 Dose-fractionation

Dose-fractionated data acquisition is achieved by repeated scanning of the same specimen area, thereby recording 4D-STEM movies. To utilise the full electron dose joint parameter estimation is first performed by averaging the power spectra and the Ronchigram cross-correlations over the dose fractions. After initialising the probe and scan grid from these common parameters, separate ptychographic reconstructions are performed for each dose-fraction as depicted exemplarily in Fig. 4 g. The resulting ptychographic movies are then subjected to established motion correction and dose-weighting routines [7, 27, 28].

### 2.5 Per-particle refinement

As schematically illustrated in Fig. 5 a, the vertical position of a protein within the ice layer may deviate from its centre or from the estimated local defocus [48, 49]. To estimate this position in beam direction ptychographic reconstructions are performed for each particle using a range of systematic offsets onto the defocus (Fig. 5 b).

**Fig. 5.**
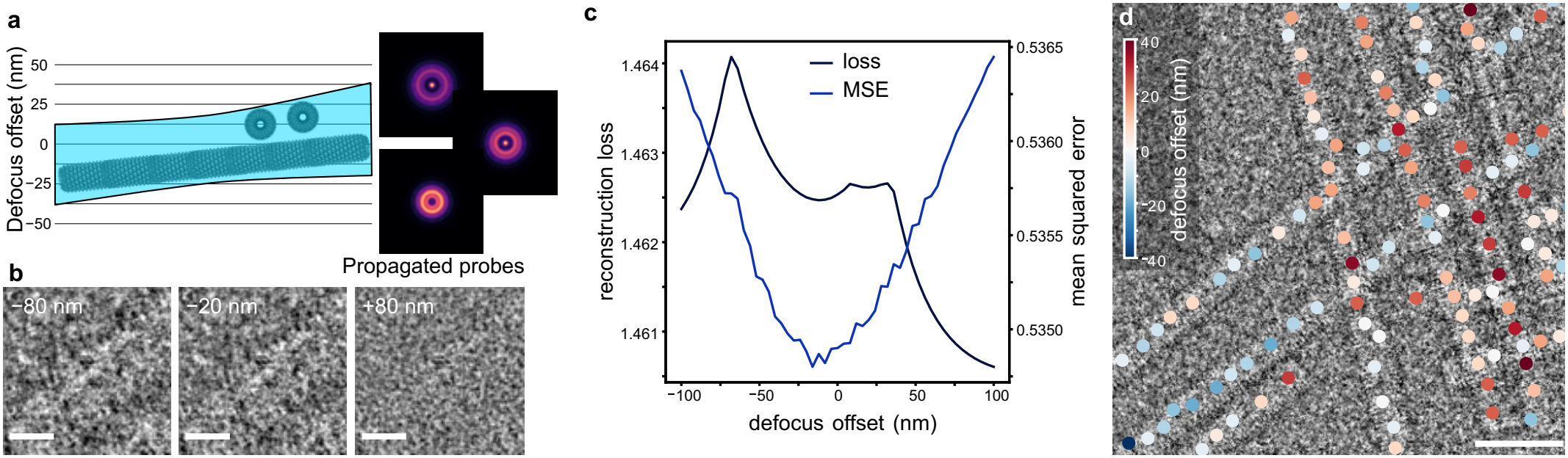
Reference based estimation of local defocus. (a) Schematic of focal series for local defocus determination. The probes are Fresnel-propagated away from the average micrograph defocus and reconstructions are performed at this fixed defocus. (b) Reconstructed TMV particle at *−*80 nm, *−*20n? m and +80n? m offset from the average defocus. (c) Reconstruction loss and mean squared error between reconstruction and projected reference for a single particle. (d) Application of the procedure to a single scan reveals TMV rods at different heights. The scalebar represents 10 nm in (b) and 50 nm in (d).

Upon visual inspection, it is difficult to ultimately judge which of the three results is best. Additionally, the loss curve in Fig. 5 c reveals several local minima. Instead we anticipate that a first three-dimensional model has been built and the optimal defocus is estimated using a reference-based approach [30]. As the 3D-map contains information from a large number of particles it can be assumed that the defocus is optimal where the agreement between a projected reference map and the reconstructed particle image is maximal. The mean square error (MSE) in Fig. 5 c reveals only a single, unique minimum which indicates that at the particle considered here, the actual defocus was approximately 20 nm lower than in the initial reconstruction. Note that this also demonstrates how volatile the blind simultaneous reconstruction of specimen and local focus can become if solely based on loss minimisation employing naturally noisy 4D-cryoSTEM data – for example, the loss is significantly lower at a focus of +80 nm where the particle could hardly be recognised in Fig. 5 b.

Mapping the minima of the MSE curve for all TMV segments in the micrograph in Fig. 5 d reveals that the method allows for the separation of particles along electron beam direction. The TMV rods oriented approximately vertically are located at a height of about +20 nm with respect to the focus of the initial reconstruction, whereas the TMV rods running diagonally lay at *−*12 nm on average, being consistent with the known TMV diameter of 18 nm. Overall, the defocus is consistent within single rods demonstrating the stability of the approach. Exceptions appear at the ends or crossings of the TMV rods, which must be expected because the projected reference will be inconsistent with the reconstructed particle.

### 2.6 Ptychographic single-particle analysis

In the following the performance of the developed workflow is first demonstrated without dose-fractionation by application to tobacco mosaic virus, horse spleen apoferritin and Hepatitis B virus (HBV) capsids (Fig. 4 a-c). Additionally, EMPIAR-12236, a 4D-STEM dataset of mouse apoferritin, was analysed (Fig. 4 d). In the second part dose-fractionation is demonstrated using TMV and the dose damage is studied. For the TMV, horse spleen apoferritin and HBV datasets an accurate calibration of the scanning geometry was performed using layer-line analysis of TMV. As a consequence the Ronchigram cross-correlation analysis was utilised for defocus determination. In the case of EMPIAR-12236 the scanning geometry was estimated for each scan separately as no such calibration was available.

#### 2.6.1 Standard SPA

##### Tobacco mosaic virus

TMV is an established test specimen with helical structure. The micrographs as depicted in Fig 4 a, representing the phase of the ptychographic reconstruction, show the TMV rods with high contrast; the characteristic layer lines are visible both in real and reciprocal space. After helical refinement, a final map resolution of 3.7 Å was obtained using 3607 particles as determined from the Fourier shell correlation plot in Fig. 4 e, compared to prior ptychographic SPA studies at 6.4 Å [25].

This resolution is confirmed by visual inspection of the map shown in Fig. 4 a. The slab view, representing a vertical section of the rods, reveals that the secondary structure of the virus is well resolved and that both the helical pitch of the alpha helices and the side-chains branching from the polypeptide backbone are clearly visible. The obtained ResLog B factor of 220 Å^2^ is consistent with the number of picked particles and the obtained resolution (see Suppl. Fig. 9).

Application of the per-particle defocus determination resulted in a map with a resolution of 3.7 Å. The lack of resolution improvement is not surprising as the deviations from the average defocus of a few tens of nanometers are small compared to the depth of focus of 221 nm at the selected semi-convergence angle of 4 mrad [50].

##### Horse spleen apoferritin

In addition to the helical TMV, the developed workflow was also applied to another well characterised cryo-EM specimen, horse spleen apoferritin. Visual inspection of the micrograph in Fig. 4 b shows the spherical particles with high contrast. After homogeneous and local refinement with octahedral symmetry a map with 4.1 Å resolution was obtained from 4999 particles. At this resolution, the map shows the helical pitch of the alpha helical regions and bulky side chains are visible.

To demonstrate the CTF-free contrast transfer of ptychography we utilise the frequency-resolved normalised correlation between the projected reference and the extracted particle patches as plotted in Fig. 4 f. The correlation plots were averaged over a complete micrograph to increase the signal to noise ratio. While the curve of the CTEM-based dataset (blue) shows clear oscillations owing to the CTF no such oscillations are observed in ptychography (black), instead a nearly monotonic decrease in correlation is found. The CTF oscillations are not present in the lowest frequencies as the CTEM image is significantly impacted by amplitude contrast there.

##### Hepatitis B and comparison with iDPC

In addition to the standard protein samples discussed above DNA-filled HBV capsids were investigated. The virus particles are approximately 32 nm in diameter, as can be seen in the ptychographic micrograph in Fig. 4 c. After refinement enforcing icosahedral symmetry a map with 4.7 Å resolution was obtained from 4049 particles.

The full map demonstrates the icosahedral capsid with the spikes extending from it. In these spikes, the pitch of the alpha helices is visible as highlighted in Suppl. Fig. 5. Since the DNA filling does not share the same symmetry as the capsid but is disordered it is not visible at the chosen threshold.

To compare 4D-STEM ptychography with integrated differential phase contrast imaging (iDPC) – another STEM-based phase contrast method – the same specimen grid was investigated using iDPC. Here the semi-convergence angle needed to be reduced to 2 mrad as iDPC micrographs recorded at 4 mrad showed very low particle contrast according to Suppl. Fig. 6 a. Utilising the same number of particles as in the ptychographic reconstruction a map with a resolution of 9.0 Å was calculated.

In this example, the ptychographic workflow demonstrates increased particle contrast at 4 mrad when compared to iDPC imaging. Importantly, ptychographic SPA is more robust against the reliability of the experimental focusing procedure which is especially important when using low-contrast carbon supports. While the contrast and resolution of iDPC imaging quickly degrades when deviating from in-focus conditions [51], the ptycho-graphic workflow is able to determine and correct for the experimental defocus.

##### Apoferritin EMPIAR-12236

The developed processing pipeline was additionally applied to EMPIAR-12236, a 4D-STEM dataset of mouse apoferritin published in [25]. Using the developed processing workflow we were able to increase the resolution of the 3D-map from the initially published 5.8 Å to 3.7 Å as shown in Fig. 4 d. At the same time the number of utilised particles was increased from 11,552 to 22,198. Analysis of the retrieved scanning geometries showed a small shear, the demonstrated increase in resolution can be at least partially attributed to its inclusion in the reconstruction (Suppl. Fig. 7). This illustrates both the robustness of the workflow and the applicability of the processing pipeline for different acquisition conditions. The better resolution of the mouse apoferritin 3D-reconstruction compared to horse spleen apoferritin 3D-reconstruction is a result of the increased number of particles and the excellent ice quality.

### 2.7 Dose-fractionated SPA of tobacco mosaic virus

Dose-fractionated acquisitions of TMV were performed utilising three dose fractions at a total dose of 23 e^−^ / Å^2^. An exemplary aligned and dose-weighted micrograph together with the reconstructed phases of the three single fractions is depicted in Fig. 4 g. The aligned and dose-weighted micrograph shows significantly higher contrast when compared both to the single fractions and the standard ptychographic micrograph in Fig. 4 a, showing the successful application of dose-fractionation. Note that this demonstrates that for doses smaller than 10 e^*−*^/ Å^2^, stable and independent reconstructions are obtained for each scan, being a result of the robustly determined probe prior to the ptychographic reconstructions.

From these micrographs an initial 3D-map with a resolution of 3.5 Å was generated (Fig. 4 i). This map was further refined using reference based motion correction and empirical dose-weighting to obtain a 3D map with a resolution of 3.3 Å utilising a total of 5330 particles, showing that dose-fractionation improves the spatial resolution of cryo-ptychography by approximately 0.4 Å in this case.

Figure 4 g additionally includes the measured local particle movements relative to the first frame as colorcoded data points. For reasons of clarity, only the lengths of the shift vectors are depicted, here amounting to significant magnitudes of up to 7 Å. Analysis of the observed shifts (including the full-frame motion) revealed a mean translational change of 5.5 Å during the acquisition [52] (Suppl. Fig. 8 a). The mean particle shift is slightly larger than previously reported CTEM-based results of 4.2 Å and 2.9 Å, determined at a similar dose of 20 e^*−*^/ Å^2^ [53]. This is not surprising as the total acquisition time of STEM movies of at least 30 s is significantly longer than that of the CTEM-based acquisitions (1 s and 4.4 s respectively) [53] with the employed detector. The largest contribution to the overall motion is given by the rigid full-frame motion. Removal of this contribution results in a much smaller mean shift of only 1.6 Å (Suppl. Fig. 8 b).

Compared to the map from single acquisitions, an overall increase in map quality is observed, as highlighted in Fig. 4 h by the visibility of the side chains. The improved 3D-reconstruction quality is confirmed by ResLog analysis where the B-factor was improved from 220 for single scans to 194 for the dose-fractionated acquisition (Suppl. Fig. 9).

From this high resolution map an atomic model of the protein part of TMV was built using ModelAngelo [31], which is displayed in Fig. 4 h. Model-building was supported by supplying the published FASTA sequence (PDB ID: 6sae) [54]. Since the TMV is disordered near the centre of the virus, no accurate model could be determined in that region. Additionally, the RNA of the TMV was not modelled by ModelAngelo as it does not build nucleic acids into cryo-EM densities. Despite that the obtained model fits the protein part of the virus very well. Inspection of the alpha helical region reveals that most side chains of the model are contained in the isosurface of the map. Here the oxygen-based negatively charged side chains are an exception since they are known to be destroyed first by beam damage [53]. Comparison of the model with a known literature model (PDB ID: 6sae) [54] resulted in an RMSD of 0.71 Å, calculated using the matchmaker tool in ChimeraX [55, 56], which additionally confirms the quality of the built model based on the determined ptychographic map.

The dose-fractionated data additionally allows to study dose damage in 4D-STEM ptychography. To this end, we utilise both 3D-reconstructions performed for each dose-fraction separately, and the layer lines appearing in the power spectra of the micrographs as displayed in Fig. 6.

**Fig. 6.**
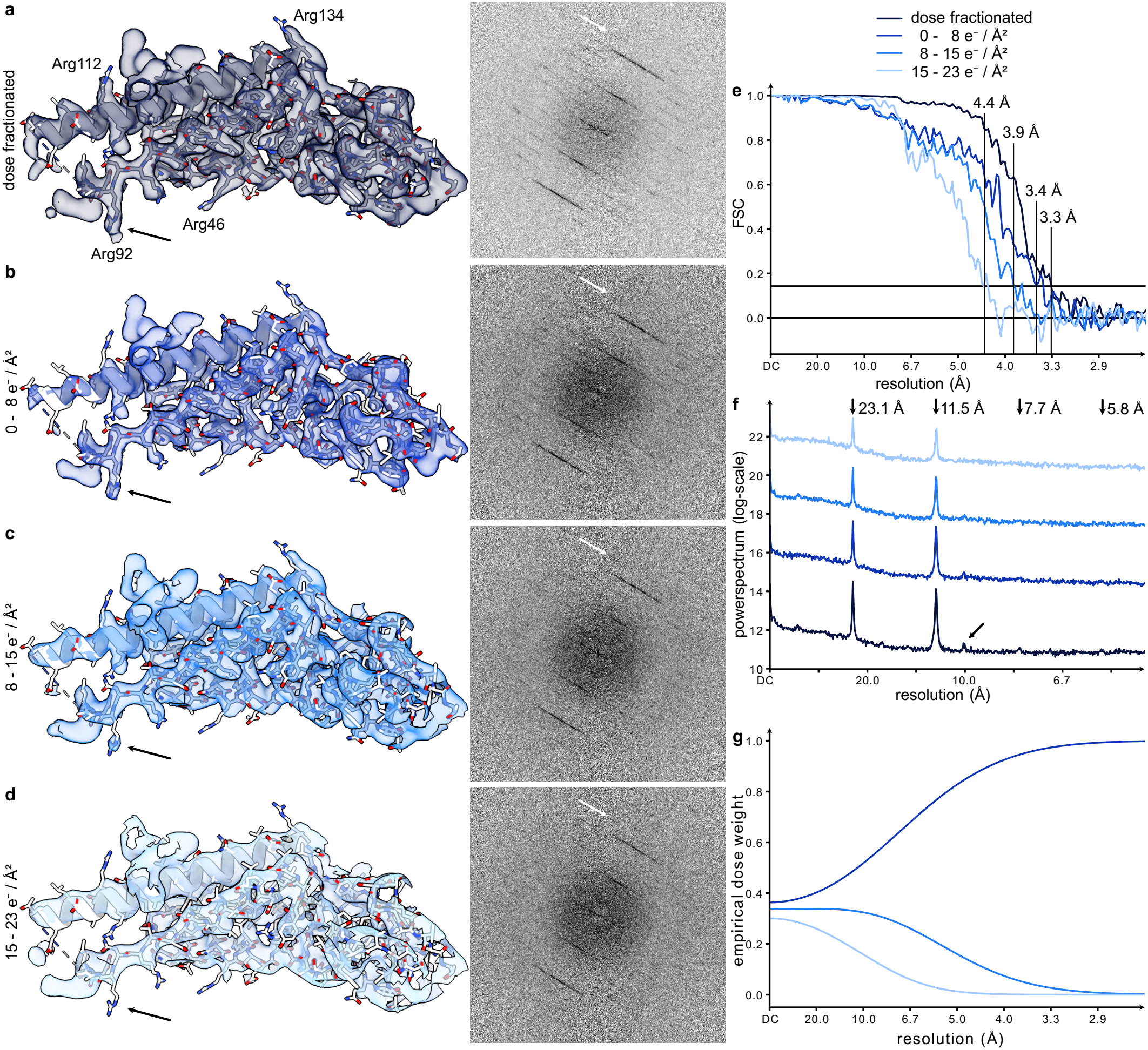
Dose damage in ptychographic SPA. Asymmetric units of TMV (3D-reconstruction) from a (a, left) dose-fractionated acquisition and (b-d, left) the three separate dose fractions of 0-8, 8-15, 15-23 Å^2^, respectively. The black arrows highlight the Arg92 side chain. The power spectra of single micrographs are shown on the right side of the same panels. (e) FSC curves of the four 3D-reconstructions supporting the reduced resolution of the single dose fractions. (f) Linescan along the power spectra in (a-d), showing the fading of the high resolution layer lines. The white arrow highlights the same 10 Å feature as in the 2D power spectra in (f). (g) Empirical dose-weights obtained from reference-based motion correction.

The layer lines are best visible in the power spectrum of the dose-fractionated micrograph in Fig. 6 a. Here, the first two layer lines and a number of secondary features can be seen. The line scan through the power spectrum in Fig. 6 f even reveals a faint third and fourth layer line at 7.7 Å and 5.8 Å respectively. While both the two layer lines and the secondary features are visible in the first dose fraction in Fig. 6 b, their contrast is reduced. The contrast of the secondary features is strongly reduced in the second dose fraction (Fig. 6 c) and completely absent in the third dose fraction in Fig. 6 d, being again confirmed by the line profiles in Fig. 6 f.

As expected [27, 57], the 3D-reconstruction of the first fraction shows the highest resolution at 3.4 Å, while the subsequent fractions result in resolutions of 3.9 Å and 4.4 Å. This trend is confirmed by both visual inspection of the maps in Fig. 6 b-d where the coverage of the sidechains is reduced by dose damage, and by the analysis of the FSC in Fig. 6 e. Importantly, the successful application of motion correction and dose-weighting are confirmed by the fact that the dose-fractionated map shows both higher resolution and strongly increased SNR throughout the whole spatial frequency spectrum compared to the single fractions. This is demonstrated by both visual inspection of the map in Fig. 6 a and the FSC in Fig. 6 e.

Analysis of the empirical dose-weights displayed in Fig. 6 g reveals that low-frequency information is retained from all fractions. In contrast, high-frequency information is only retrieved from the first fraction with the second one not contributing beyond 4.0 Å, and the third dose fraction not providing information beyond 6.7 Å.

## 3 Discussion

Using the workflow developed in this study we successfully demonstrated ptychographic single-particle analysis down to 3.3 Å resolution for dose-fractionated 4D-STEM cryo-ptychography, and 3.7 Å for unfractionated acquisitions. This resolution enabled the automatic building of an atomic model of TMV using ModelAngelo [31]. A similar resolution has been previously obtained using iDPC [18], a non-iterative approach approximating center-of-mass imaging [51, 58]. With this workflow, we built a robust framework for ptychographic image reconstruction while miminzing human intervention. This workflow largely relies on automated ptychographic processing from low dose data, which are followed by established single-particle processing workflows.

The demonstration of dose-fractionation additionally opens up new opportunities for future acquisitions. If higher resolution is desired more and finer dose fractions need to be acquired to limit dose damage. To increase the general particle contrast more dose fractions at higher total electron doses should be acquired. Importantly this can be done without impacting resolution owing to the dose-weighting. Here, the recent introduction of new detector hardware such as Timepix3/4 [38, 39, 59, 60] with even further increased speed and number of pixels will additionally allow for both faster acquisitions and the application of larger defoci as well as SCAs. While higher SCAs can be used to increase the maximum achievable resolution, larger defoci will allow for larger scan steps, resulting in higher doses per diffraction pattern and, as a consequence, open up the way for finer dose-fractionation. In addition to the SCA, ptychographic SPA is expected to profit from high coherence, e.g., via cold field emission guns [61] potentially combined with corrected electron optics [62, 63].

The resolution obtained using the dose-fractionated TMV dataset (3.3 Å) is equivalent to first dosefractionated CTEM-based experiments performed on the same virus (3.4 Å) [53]. Further comparison is hindered by differences in the utilised acquisition parameters. While the ptychographic dataset contains a larger number of particles, it was also recorded using coarser dose-fractionation and at a lower total dose compared to the CTEM-based dataset. It is worth noting that the resolution of CTEM-based imaging of TMV has since been increased to below 2 Å by utilising improvements in hard-ware and image processing [54]. Similar improvements in STEM will likely require finer dose-fractionation, larger semi-convergence angles and more precise scanning systems because counting noise puts systematic constraints to the attainable precision in post-acquisition scan position refinement [40]. From the current perspective, it can be anticipated that the ptychographic SPA workflow devised here solved inherent challenges of low-dose cryo-ptychography and is thus generic and, consequently, applicable to 4D-cryoSTEM experiments taken with improved hardware and/or largely different acquisition parameters. In particular, the workflow is as well directly applicable to weakly scattering specimens in materials science, e.g., for imaging organic frameworks [64] or perovskite nanostructures [65].

Application of the workflow towards ptychotomographic reconstructions of thick specimens based on 4D-STEM suggests itself as an additional research line to pursue [66]. In that respect, the robust local defocus estimation reported here can be seen as a pre-requisite for the reconstruction involving high tilt angles. Furthermore, the thicker specimens used in tomography will benefit most from physically more reliable interaction models [45] that can straight forwardly be used within our ptychographic workflow, such as the inverse multislice scheme [67–69].

In summary, we presented a workflow rendering ptychographic SPA a viable tool in structural biology with competitive spatial resolution. While the developed implementation utilising python programs allows for the straight forward execution on remote servers or clusters, an implementation supporting graphical user interfaces is of interest as it will allow for easier inspection of the data and manual exposure curation.

## 4 Acknowledgement

The authors acknowledge funding from the European Research Council within the Horizon Europe innovation funding programme under Grant Agreement 101118656 (ERC Synergy project 4D-BioSTEM). A.K.S. acknowledges support from the German Research Foundation (DFG) through the Emmy Noether program (project number 394455587) and TRR179 (project number 272983813). We thank Tillman Schäfer and Daniel Bollschweiler (Max-Planck-Institut fuer Biochemie, Martinsried) for preparing the grids of the HBV capsids. Support from the team of Quantum Detectors, Inc. (UK) for the installation and read-out of the Timepix4 detector at Ludwig-Maximilians-University (LMU) Munich is gratefully acknowledged. The authors gratefully acknowledge the computing time granted by the JARA Vergabegremium and provided on the JARA Partition part of the supercomputer JURECA at Forschungszentrum Jülich [70].

## 5 Author contributions

T.L. designed the processing workflow, implemented it in the TorchSlice-package and performed the data processing. D.M. and T.L. collected the ptychographic SPA datasets and performed the SPA processing. D.M. initiated the dose-fractionation approach. J.Z. proposed the rCTF, provided initial implementations of Ronchigram cross-correlation and iterative bright field movement estimation, and provided the implementation for the projection of reference volumes. F.vH. and M.S. prepared the Hepatitis B specimen under supervision of A.K.S. V.K. recorded the Apoferritin data in Fig. 2 and Fig. 3 under supervision of H.S.. T.L., A.J., M.L.L. and B.D. developed the TorchSlice package. T.L. recorded the data of the cross-grating in Fig. 2. T.L. and K.M.-C. wrote the manuscript with input from all authors. K.M.-C., C.S. and H.S. acquired funding.

## 6 Methods

### 6.1 Specimen preparation

#### 6.1.1 Tobacco mosaic virus

Tobacco mosaic virus was purified as described earlier [54]. Grids were prepared by applying 3.6 µl of solution onto a glow-discharged amorphous nickel titanium grid and plunge freezing using a TFS Vitrobot (4^*°*^C, 95 % humidity, force *−*5, 6 s blot time).

#### 6.1.2 Apoferritin

Commercial horse spleen apoferritin (Sigma-Aldrich, A3660-5X1VL) was prepared by buffer exchange of 600 µl protein solution with 1200 µl buffer solution (30 mM HEPES pH 7.5, 150 mM NaCl) using a 100 kDa spin concentrator for three hours. Using 3x Bradford reagent the concentration was determined to be 10.2 mg/ml.

Grids were prepared by applying 3.6 µl of solution onto a glow-discharged amorphous nickel titanium grid and plunge freezing using a TFS Vitrobot (4^*°*^C, 95 % humidity, force *−*5, 4.5 s blot time).

#### 6.1.3 Hepatitis B virus capsids

##### Cp183 expression and purification

All chemicals were purchased from Merck or Carl Roth unless otherwise stated. Full-length Hepatitis B virus genotype D core protein, comprising residues 1-183 (Cp183), was expressed in E. coli BL21(DE3) from a pRSF1b vector using a tagless, codon-optimized insert. Cells were grown in LB medium at 37 °C, induced with IPTG, and incubated overnight at 20 °C.

Harvested cell pellets were resuspended in a lysis buffer (50 mM Tris pH 8.0, 300 mM NaCl, 5 mM EDTA and 10 mM MgCl2) supplemented with protease inhibitor mix (SERVA), 0.2 % IGEPAL CA-650 and DNase I (SERVA). Cells were disrupted by sonication, and the lysate was clarified by centrifugation. The supernatant was loaded onto a 10–60 % sucrose step gradient prepared in a buffer containing 50 mM Tris pH 7.6, 300 mM NaCl, 5 mM EDTA and 5 mM DTT and subsequently ultracentrifuged overnight at 160,000 g. Cp183-containing fractions were pooled and diluted twofold with HBS-300 buffer (150 mM HEPES pH 7.5, 300 mM NaCl). Capsid protein was then precipitated by stepwise addition of solid (NH4)2SO4. After centrifugation at 30,000 g for 45 min, pellets were resuspended in 20 mL buffer (50 mM HEPES pH 7.6, 150 mM NaCl, 5 mM EDTA, 5 % sucrose) and dialyzed overnight into the HBS-300 buffer. After a final centrifugation step (30,000 g, 20 min) capsids were concentrated, supplemented with 5 % glycerol and flash frozen until further use.

##### Capsid disassembly and RNA removal

Cp183 capsids were dialyzed overnight into disassembly buffer (50 mM HEPES pH 7.5, 1.5 M guanidine, 0.5 M LiCl, 2 mM DTT). The precipitated RNA was then sedimented via centrifugation at 30,000 g for 15 min. The disassembled dimers were then separated from capsids via size exclusion chromatography (HiLoad 16/600 Superdex 200 pg), equilibrated with the same buffer. Dimer fractions were combined for subsequent steps.

##### Assembly of dsDNA capsids

The in vitro packaging of nucleic acids into Cp183 capsids has been previously detailed. Briefly, as a stand-in for the viral relaxed circular DNA (rcDNA), the similarly sized (2686 bp) bacterial plasmid pUC18 was amplified in E. coli DH5αcells and purified using the PureLink HiPure Expi Plasmid Gigaprep Kit (Invitrogen) according to the manufacturer’s instructions.

Pooled dimer fractions were combined with a two-fold molar excess of pUC18, equating to a single plasmid for 120 dimers or one T4 capsid. The solution was diluted threefold with water to a final LiCl concentration of 0.5 M, thus initiating capsid reassembly, and incubated at room temperature for 1 h. The dsDNA-filled capsids were separated from remaining pUC18 via sucrose gradient centrifugation following the procedure described above. Successful encapsidation was verified by native agarose gel electrophoresis (NAGE) to confirm the co-localization of nucleic acids and proteins via intercalating DNA stain (SERVA) and Coomassie staining, respectively. Finally, gradient fractions containing dsDNA capsids were pooled and dialyzed into the final buffer (50 mM HEPES pH 7.5, 150 mM NaCl, 5 mM EDTA).

##### Preparation of the specimen grid

4 µl of a 1 mg/ml solution of HBV capsids were added to a glow-discharged Quantifoil holey carbon grid and plunge frozen using a TFS Vitrobot (4^*°*^C, 95 % humidity, force 5, 3 s blot time).

### 6.2 Data acquisition

single-particle data was recorded on a Titan Krios G4 (Thermo Fisher Scientific) operated at 300 keV with an SCA of 4 mrad. The HAADF detector was used for navigation. Focusing was performed using both the HAADF and the Ronchigram. A Dectris Arina detector was operated in binned mode at a frequency of 100 kHz, such that a single scan of 1024 *×* 1024 STEM scan points was recorded in 10–12 s with a scan step of 5 Å. Dosefractionated data was acquired by reducing the beam current and recording multiple scans at the same specimen location. Detailed acquisition parameters for the datasets used to produce the figures appearing in the manuscript are summarised in Suppl. Table 1.

### 6.3 TorchSlice framework

Data processing was performed within the TorchSlice framework [24, 71]. In addition to its core functionality being multislice simulations and inverse multislice ptychography, TorchSlice comprises a large number of 4D-STEM techniques such as STEM imaging *via* virtual detectors [72], single side band ptychography [21], Wigner distribution deconvolution ptychography [42], parallax [19, 32] and shadow montage imaging [20], together with the methods described in the present work. The TorchSlice framework makes full use of the automatic differentiation framework of PyTorch [41], thereby allowing the probe and specimen to be parameterized and optimized directly in terms of physical parameters, such as aberration coefficients. All routines utilise the same basic parameter objects, so that the same aberration object returned, e.g., by single-sideband ptychography, can be used for parallax imaging or partially coherent multislice ptychography.

To facilitate processing of the single-particle data, the routines are separated into a set of programs, each performing a single task, such as the calculation or evaluation of the Ronchigram power spectra. Information is stored in a single YAML file per scan, allowing it to be easily readable by both humans and machines [73]. In the YAML files, compact evaluation results and storage paths to extensive or intermediate results are saved. This allows for separation of the large experimental 4D-STEM data, e.g., on a fast SSD, from the results, e.g., on a larger hard drive with a different backup schedule. Separating tasks into these programs also enables straight forward calculations across different computer systems or parallel processing of different tasks on the same system.

### 6.4 Estimation of the rCTF

Defocus estimation using the Ronchigram contrast function (rCTF) consists of two steps, the calculation of the power spectrum of the shadow image, followed by the determination of the rCTF parameters. In the presence of defocus and astigmatism, the magnification of the shadow image is proportional to the inverse of the defocus-astigmatism matrix **H**

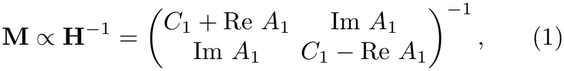

which is constructed from the defocus *C*_1_ and the complex-valued astigmatism *A*_1_ [33, 74]. The rCTF for the observed Ronchigrams is given by [34]

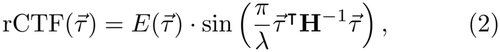

where 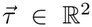 is the Fourier-conjugate coordinate to the scattering vector 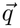 inside the Ronchigram and *λ* is the electron wavelength. The function 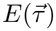 represents damping envelopes due to, e.g., partial coherence. We evaluate the power spectra and thus the squared rCTF, such that its zero crossings occur as minima.

A difficulty arises because the rCTF governs only the transfer of spatial frequencies within the Ronchigram. When Fourier-transforming the diffraction patterns, the aperture dominates the raw power spectrum in that it causes oscillations with nearly constant, and typically higher frequency than those of the rCTF. Thus, special care is needed to suppress the influence of the aperture; for this reason, the mean CBED is subtracted from the individual ones. The mean-subtracted CBEDs are then zero-padded to at least twice the size of the detector, and the squared moduli of their Fourier transforms are accumulated. In some situations, artifacts of the aperture remain, e.g. if the evaluated area varies in thickness, the mean CBED can be an insufficient representation of the diffraction patterns. Here, a Gaussian weighting mask centred on the optic axis can optionally be applied in addition to suppress these effects. A standard deviation of 0.5 bright field radii has proven suitable. Similar effects arise when prominent bright field movement is present. Here, the same weighting filter can be applied, or the evaluation can be performed after correction of the bright field movement, i.e. with CBEDs interpolated such that the Ronchigrams are centred around the optic axis.

Since the defocus-astigmatism-matrix is symmetric, its inverse is also symmetric. Consequently, only three unknowns must be determined (the two diagonal elements and one off-diagonal element). During optimisation, the rCTF is parameterised by this inverse matrix and the real defocus and astigmatism are reported only at the end. Using this formulation, the determination of the rCTF is akin to the CTF determination problem in CTEM [35, 75].

During the rCTF determination, first a background-subtracted amplitude spectrum is calculated, and a starting value for the defocus is obtained by an exhaustive search in a given range. The defocus and astigmatism are then refined using gradient-based optimisation. Second, gradient-based optimisation of the envelope, background, defocus and astigmatism is employed to minimise the mean squared error between the modelled and experimental power spectra. While the inclusion of the parameterised envelope and background significantly increases the agreement between model and experimental data, it has little effect on the determined parameters, as shown in Suppl. Fig. 11.

The background-subtracted square root of the power spectrum will be denoted 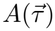 in the following, where 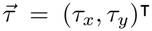 is the Fourier conjugate variable to the spatial frequency 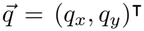. Background estimation is performed by real-space convolution with a square box. Since the centre of the power spectrum is dominated by a strong peak which can distort the background estimation, it is replaced with the mean of the surrounding area. A radius of 10 pixels has proven sufficient in our study. The correct rCTF parameters are then determined by maximising the normalised correlation between the background-subtracted amplitude spectrum and the modulus of the rCTF,

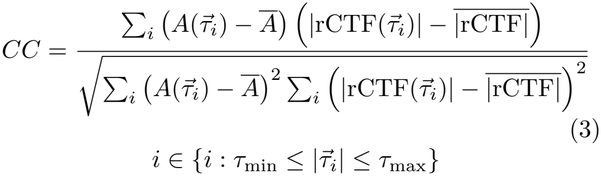

where the bar denotes the mean value. This correlation coefficient is calculated in a user-selected range between *τ*_min_ and *τ*_max_. The rCTF is given by eq. (2) using a flat envelope.

First, a rough estimate of **H**^*−*1^ is obtained using an exhaustive search, followed by gradient-based refinement of these starting parameters. The correlation coefficient is evaluated at a range of (inverse) defoci between 1*/*Δ*f*_max_ and 1*/*Δ*f*_min_ while assuming the astigmatism to be zero. This step is performed on azimuthally averaged data to reduce computational cost. Starting from these initial parameters, the correlation coefficient of the 2D-amplitude spectrum and the 2D-rCTF model is maximised using limited-memory Broyden–Fletcher–Goldfarb–Shanno algorithm (L-BFGS) [76].

To model the full power spectrum 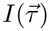, the background and envelope are modelled using exponentials of polynomials with user-defined degree. These are initialised using the estimated defocus and astigmatism. A radially symmetric background and envelope are used. All parameters (background coefficients *θ*^BG^, envelope coefficients *θ*^env^ and defocus-astigmatism matrix *H*) are then refined further by minimising the mean squared error between the background-subtracted power spectrum and the squared rCTF multiplied by the envelope,

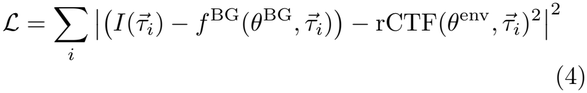

where the summation is again restricted to the user-defined fit range. If required, an additional regularisation term based on the rectified linear unit (ReLu) [77] can be added

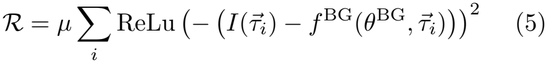

which enforces that the background-subtracted power spectrum is positive. A regularisation strength *µ* between 0 (no additional regularisation) and 0.1 has proven useful.

After rCTF determination, the final correlation coefficient and two diagnostic figures are reported utilising either the background-corrected amplitude spectrum or the power spectrum. The first figure is a plot of the 2D-rCTF model overlayed atop either the background-subtracted amplitude spectrum or the background-subtracted full power spectrum. Secondly, a radial average of the spectrum (blue) and the rCTF model (red) is reported (see Fig. 2), along with the normalised correlation coefficient calculated between the rCTF maxima (black). This correlation coefficient is calculated on the full two-dimensional spectra. In the presence of astigmatism, all radial averages are performed along elliptical iso-curves of the rCTF [37]. The resolution of the fit is reported at the point where the normalised correlation coefficient passes below a threshold of 0.25. While this threshold is arbitrary, a higher resolution at the same threshold clearly indicates a higher fit quality.

To extend the rCTF determination for calculating the local defocus, the power spectra are calculated in patches of probe positions. The defocus variations are parameterised using a grid of cubic B-splines [35]. This parameterisation ensures smoothness of the fitted defocus landscape, while the degree of the defocus variation can be configured using the number of control points.

Instead of evaluating all patches separately and fitting a surface to the resulting defoci, we treat all patches simultaneously and optimise the parameters of a spline grid describing the underlying defocus variation. In this way, a constant astigmatism across the scan area can be guaranteed, and robustness against low-contrast patches is enhanced.

The local defocus landscape is initialised to a global estimate determined from the average power spectrum, and the optimisation goal is the sum of correlation coefficients for each patch

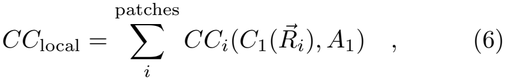

where the local defocus 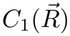 is evaluated at the centre 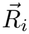of each patch. Importantly, the parameterisation is performed using the real defocus and astigmatism rather than the entries of the inverse defocus-astigmatism matrix. This change is required because only defocus is allowed to vary across the field of view, while other parameters, such as astigmatism, are assumed to be constant. The optimisation of the spline coefficients and the global astigmatism is performed using L-BFGS [76]. This optimisation is performed using the background-subtracted amplitude spectra without inclusion of an envelope to reduce the number of parameters to be optimised. After optimisation, a local defocus-adjusted power spectrum is reported by rescaling the patch-power-spectra according to

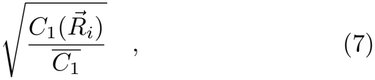

which aligns the ring patterns onto each other [35]. If the local defocus refinement is successful, more rings are visible, and the reported fit resolution is increased.

### 6.5 Ronchigram cross-correlation

The properties of the Cartesian scanning grid can be represented by a 2 *×* 2 matrix **R** encoding the real-space probe shift resulting from a step in the fast and slow scan directions. Since the magnification of the shadow image is known, the movement in the shadow image 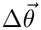 can be described (in mrad units) by

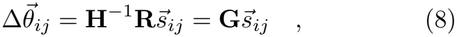

where 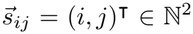 represents the number of steps in the fast and slow scan direction.

To determine the scan coordinates, the average cross-correlation between neighbouring shadow images is first calculated for up to *n*-neighbours in the fast and slow scanning direction. During calculation of these cross-correlations, the mean-subtracted CBEDs are zeropadded to twice the size of the detector, followed by the calculation and accumulation of the cross-correlations.

A naïve approach of detecting the maximum of each cross-correlation will likely fail in cases with a low SNR, or in situations where the true shadow step is larger than the shadow image. Instead, the matrix **G** is estimated directly by maximising the sum of the cross correlations sampled at the points predicted by these shadow steps

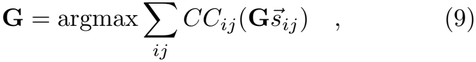

where *CC*_*ij*_ denotes the mean cross-correlation between neighbours separated by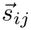 scan steps.

The optimal **G** is determined by a two-step procedure. First an exhaustive search with user-defined step sizes is performed, typically utilising a step size between one and four times the detector pixel size. This exhaustive search is further restricted by user-defined limits on the minimum and maximum length and for the shear and relative length difference between the two columns of **G**. The maximum length is usually chosen as twice the bright field radius, while the limits on relative shear and length difference are set to 0.1. This appears reasonable since the microscope is typically aligned to minimise astigmatism and to ensure that the scanning geometry is square and symmetric. This initial estimate is then further refined by gradient-based optimisation of **G** utilising the L-BFGS algorithm [76]. Nearest-neighbour interpolation is used during the exhaustive search, whereas during gradient-based refinement, interpolation is performed by analytically evaluating the inverse Fourier transform of the cross-correlation to ensure smoothness of the gradients [78].

If the defocus-astigmatism matrix is known from the rCTF analysis described above, the full matrix **R**

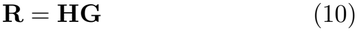

including shear and a difference in scan step in the slow and fast direction, can be determined. If the scanning geometry is known, the defocus, astigmatism, and the orientation of the scan with respect to the camera can be obtained:

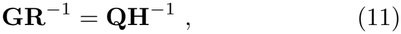

the rotation matrix **Q** and **H**^*−*1^ can then be separated using polar decomposition, which reduces a general matrix into the product of a unitary matrix (representing the orientation between the scan and camera coordinate systems) and a symmetric matrix representing the inverse defocus-astigmatism matrix.

In the presence of local defocus variations, the cross-correlations are calculated in patches of scanning positions. After the determination of the local defocus, these are rescaled according to

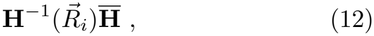

which is the ratio of the local magnification of the Ronchigram at 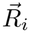 to the mean magnification. If a significant local defocus variation is present, the cross-correlation peaks should appear more symmetrical and sharper, indicating a successful recovery of the local defocus. Note that the rescaling procedure is incompatible with the extraction of the power spectrum from the peak shapes of the Ronchigram cross-correlation due to the different rescalings of the power spectra and the cross-correlations.

The patch cross-correlations can also be used to determine the local defocus. In this case, the local defocus parameters and the global astigmatism are determined by maximising the sum of the cross-correlations for all patches sampled at the points predicted by the current defocus model. In this case, a constant scanning geometry is assumed.

The modulation of the cross-correlation peaks can be understood in the following way. Let 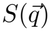 and 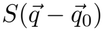 be two shadow images inside the Ronchigrams of two neighbouring scan positions with sufficient probe overlap. Then, the inverse Fourier transform of the power spectrum of *S* appears centered around the cross-correlation peak, being the auto-correlation in case the two Ronchigrams are identical except for the shift of their internal features. In practice, this is only approximately the case, because the illuminated regions differ slightly. Importantly, this opens up an alternative method for obtaining the power spectrum of the shadow image and thus to gauge the defocus [33]. When estimating the power spectrum of the shadow image using the peak shape of the cross-correlation peaks, all cross-correlation peaks are shifted to the origin using the predicted shifts. The power spectrum is then obtained from the real part of the inverse Fourier transform and processed using the same rCTF analysis as described above.

### 6.6 Bright field movement estimation

The movement of the disk, when scanning, is described using a set of two Zernike polymonials for *q*_*x*_ and *q*_*y*_, respectively, in dependence of the probe position coordinate 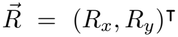. If the descan parameters are assumed constant over a set of scans, the coefficients of the Zernike polynomials are estimated by a weighted least-squares fit to the mean centre of mass at each scan position. The weights equal the variance of the centre of mass at each scan position.

The second method, applicable to single scans, consists of an inner and an outer loop. First, the cross-correlations between each CBED and the average CBED are calculated, the coefficients **c** of the polynomials 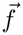 are then optimised by maximising the sum of the cross-correlations *CC* sampled at the shifts given by the polynomial

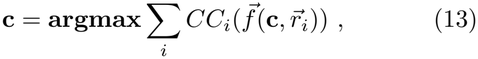

where the sum extends over all CBEDs. This inner optimisation loop is performed using the L-BFGS algorithm [76]. Using the obtained coefficients, a new aligned PACBED is calculated, and the inner optimisation loop is performed again to further optimise the coefficients. Aligning the CBEDs onto the optic axis is done in Fourier space. This outer loop is repeated until the polynomial parameters converge, typically within two or three iterations.

Importantly, the aligned PACBED and the cross-correlations need to be calculated only once per outer optimisation loop, while the inner loop consists only of evaluating the polynomial and accessing it at the positions given by its coefficients.

### 6.7 Masking of support

During preprocessing, masks are created that discard all scan points, which do not contain thin amorphous ice. To create these masks, a histogram of detector counts per scan point is first generated for all recorded datasets. The histogram is then fitted with a Gaussian mixture model using a user-defined number of components (typically 2). Each scan point is assigned to the appropriate component, and a Gaussian low-pass filter with a standard deviation of 5 pixels is applied. The final mask is obtained after thresholding at 0.5 as shown in Suppl. Fig. 10.

The component with the lower mean value is typically associated with thick support films or large amounts of contamination. Masked scan points are removed from preprocessing and not included in the ptychographic reconstructions.

### 6.8 Ptychography

Ptychographic reconstructions were performed using a singleslice object and a fully coherent object in the TorchSlice package. As the loss function, the mean-square amplitude error was chosen due to its favourable properties during optimisation of low-dose data. Stochastic gradient descent was performed with a minibatch size of 32 for a fixed number of iterations (100), where it was observed that the loss did not improve significantly. During the reconstruction, the pixelated probe and object were optimised, while the scanning grid and bright field movement were kept fixed.

Reconstructions were performed using a window twice the size of the diffraction camera, and the central part of the simulated CBEDs was used to calculate the loss. To minimise the required GPU memory, the CBEDs are stored in uint8-format and the gain or flat field reference is only applied when calculating the loss. Consequently, less than 4 GB of VRAM is required when reconstructing a subset of 512 by 512 scan positions, allowing the utilisation of consumer-grade GPUs.

Learning rate optimisation using hyperparameter tuning is performed in TorchSlice by utilising optuna with a tree-based Parzen estimator and a Hyperband pruner [46, 79, 80]. Here, the learning rates were optimised to minimise the loss after 25 iterations of ptychography.

During the reconstruction, we treat the tilt 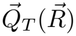 induced by the bright field movement at a probe position 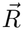 in first order by applying a phase wedge to the scanning probe in real space before interaction with the specimen, assuming that the effect of aberrations is invariant with respect to the scan position. The probe wave function at scan point 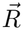 then reads

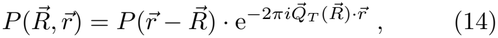

with 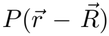 the probe calculated the conventional way, either parameterised as a pixelated array or as the inverse Fourier transform of the wave in the back-focal plane of the probe-forming lens, in which aberrations such as defocus, astigmatism and spherical aberration are accounted for by a phase plate.

For large scans, we implemented splitting a single scan into multiple patch-wise reconstructions to limit the extent and number of pixels of the resulting object. Here, each 1024 *×* 1024 scan was split into four 512 *×* 512 scans, resulting in four ptychographic reconstructions of slightly less than 4k*×* 4k pixels, with the exact number depending on the detector sampling and scan grid.

### 6.9 Single-particle analysis

All single-particle analyses were performed using cryoSPARC [7]. As input, the phase of the ptychographic reconstructions in MRC-format was used [81]. Areas which were not reconstructed (i.e. padding at the edge of the reconstruction or masked areas) were filled with Gaussian noise of the same mean and variance to match the statistics of the reconstructed area. This was required as otherwise, no particles were picked in many micrographs. For dose-fractionated acquisitions, each dose fraction is treated as a single frame in a movie, and patch motion correction is performed prior to the other processing steps. After obtaining an initial 3D model, the motion was further refined using reference-based motion correction [82]. Further steps performed during reconstruction are discussed in the dedicated sections.

### 6.10 Per-particle refinement

When performing the per-particle refinements of the ptychographic reconstructions, the particle locations and poses are first extracted from cryoSPARC. Scan positions around each of the particles are selected, allowing for ptychographic reconstructions of individual particles separately. The selection radius is chosen such that the reconstructed area is slightly larger than the extracted particle patches. Offsets are applied to the scan positions so that the projected reference and the cut ptychographic reconstruction match. For each particle, a focal series is calculated in a user-defined range (here *±*100 nm in 51 steps) by Fresnel propagating the probe from the original reconstruction by a defocus offset Δ*C*_1_ and performing a set number of ptychographic iterations (here 20). During the per-particle refinement, the probe itself is not further optimised, as the low number of electrons contributing to a single-particle can degrade the overall probe quality.

The focal series is evaluated by two metrics, the reconstruction loss and a weighted mean square error (MSE)

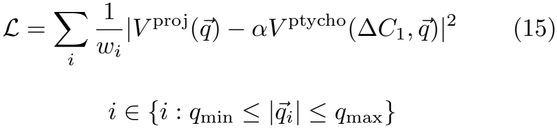

between the projected reference 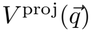 and the reconstruction 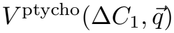 in Fourier space. The MSE is calculated over the frequency range from a minimum spatial frequency *q*_min_ (here 30 Å) to the resolution of the map. The radially symmetric weights *w* are chosen to match the SPA noise model (here obtained from the diagnostic curves plotted in cryoSPARC). Alternatively, a noise model can be approximated by the azimuthally averaged power spectrum of the particles. The scale constant *α* compensates for global scaling of the phase and is chosen by minimising the MSE in eq. (15). The scaling is applied to the ptychographic reconstruction as the MSE can otherwise be skewed by reconstructions with a very low total phase shift, e.g. at a large offset from the true defocus. Before Fourier transforming, the ptychographic reconstructions are cut to match the original particle box, and both the reference maps and the reconstruction are padded to twice their original size to reduce interpolation artefacts in Fourier space. Gold-standard single-particle refinement is enforced by using the half-map constructed from the same half-set to which the particle belongs as the reference.

The obtained MSE curves can be used to identify the per-particle defocus, and the reconstructions at this defocus can be exported for further processing. In the case of dose-fractionated acquisitions, each fraction is reconstructed separately and averaged using the dose-weights and shifts calculated during reference-based motion correction.

### 6.11 Analysis of single-particle data

#### 6.11.1 Tobacco mosaic virus

A total of 30 scans was acquired and split into 103 micrographs after discarding patches containing only support. The foci were determined to be in the ranges *−*190 and *−*800 nm and 270 and 1200 nm. From four manually picked micrographs, a 2D classification template was generated and used for further template picking. 32,386 candidate particles were identified and extracted with a box size of 512. Using 2D-classification, 4 classes consisting of 8,179 particles were selected for further processing. After two rounds of helical refinement (maximum symmetry order 140), a map with 3.7 Å resolution was obtained from 5,683 particles. Using the “Subset particles” job to split the particles into two subsets based on their per-particle scale value, 3,607 particles with high per-particle scales were identified. From these particles, a map of 3.7 Å resolution was obtained using a helical refinement job starting with a cylindrical initialisation. The initial voxel size was corrected by matching the found helical pitch of 1.42 Å to the known helical pitch of 1.40 Å, and the resolution at this corrected voxel size is reported [54].

#### 6.11.2 Apoferritin

80 scans were recorded with defoci ranging between *−*220*−*– 920 nm and 200–660 nm. From these, 285 micrographs were reconstructed. Due to the strong bright field movement present in the data, the bright field movement was determined per-scan for this dataset. Templates were generated by manually picking 50 particles, followed by 2D-classification using 5 classes. Employing the generated templates, 467,120 initial candidates were identified, out of which 62,387 particles were selected for further processing. Particles were extracted at a box size of 256 pixels, cropped to 128 pixels in Fourier space. After two rounds of 2D-classification, 12,806 particles were selected for 3D-refinement.

Homogeneous refinement (O symmetry) starting from a literature map (EMD-4905) lowpass filtered to 12 Å lead to a first 3D-reconstruction with 6.5 Å resolution. The relatively weak low-pass filtering was required due to the high symmetry of the apoferritin, which resembles a ball at lower resolutions, preventing alignment. Using Local refinement, a resolution of 6.0 Å was obtained. 3D classification was performed with three classes at a filter resolution of 6 Å and the highest resolved class was used for further refinement. The largest class containing 4999 particles was selected for further processing. Using this class, a resolution of 4.2 Å was obtained by homogeneous refinement, which was further improved to 4.1 Å using local refinement.

#### 6.11.3 Hepatitis B virus capsids

After recording 109 scans, 436 micrographs were obtained with defoci between *−*250*−*– 840 nm and 380– 940 nm. Particle picking was performed using a ring blob with an inner diameter of 15 nm and an outer diameter of 30 nm. From the initially detected 84,545 candidates 53,414 particles were extracted at a box size of 800 pixels. After 2D-classification with 50 classes, a single class containing 4052 particles was selected for further processing and downsampled to 400 pixels. The 3D reconstruction was initialised with a map of a different HBV capsid, lowpass filtered to 30 Å. Following homogenous and local refinement with icosahedral symmetry, a final map resolution of 4.5 Å was obtained from 4049 particles. The voxel size was corrected by comparison with a published atomic model of the HBV capsid (PDB ID: 6HTX) [83] and the resolution at this voxel size is reported.

#### 6.11.4 Mouse Apoferritin - EMPIAR-12236

The scanning grid was initialised using the Ronchigram cross-correlation technique, while the probe shape was determined by rCTF determination using the power spectrum extracted from the Ronchigram cross-correlation peaks. Here the amplitude-based rCTF determination without inclusion of an envelope was used. The bright field movement was determined for each scan using the iterative method.

Particle picking was performed using a ring blob with an inner and outer radius of 80 and 100 Å. The 44,000 initial candidates were extracted at a box size of 384 pixels. After two rounds of 2D-classification, 22,420 particles were selected for 3D-processing. An initial model was built by performing an *ab-initio* 3D-reconstruction with two classes while enforcing octahedral symmetry (initial resolution 20 Å). The better-resolved class was used for further refinement with an initial low-pass filter of 12 Å. After homogeneous refinement (22,198 particles, 4.0 Å) and local refinement a resolution of 3.7 Å was obtained from 22,198 particles. By comparing the reconstructed map with a literature model (PDB ID: 8RQB) [25] the actual voxel size was determined to be 0.730 Å (initial: 0.747 Å) and the resolution at this voxel size is reported.

#### 6.11.5 Dose-fractionated TMV

Data was acquired at fifteen specimen locations each by scanning three times with a defocus between *−*260 and *−*530 nm. From these scans, 55 ptychographic movies were obtained. After patch motion correction, particles were picked using templates generated from a prior reconstruction of TMV. Particle picking was performed on an initial ptychographic reconstruction neglecting astigmatism, and the particle picks were transferred to a reconstruction that included astigmatism. 24,180 candidate particles were extracted at a box size of 512, after 2D-classification 5,622 particles were chosen for further 3D-processing. Using helical refinement with a maximum symmetry order of 140, an initial map at 3.5 Å resolution was obtained and used for reference-based motion correction. Using the motion-corrected particles, a final 3D map with a resolution of 3.3 Å was obtained. The voxel size was again corrected by matching the measured helical pitch of 1.42 Å to the known helical pitch of 1.4 Å, and the resolutions at this pixel size are reported [54].

Model building was performed using ModelAngelo, utilising a map after local filtering [31].

## Data availability

The EMDB accession numbers for cryo-EM maps and PDB models (where applicable) of TMV, apoferritin, Hepatitis B virus capsids have been deposited as EMD/PDB IDs: EMD-100046/PDB-34NO, EMD-100057/PDB-34OI, EMD-100042, EMD-100078/PDB-34OZ, EMD-100063.

## References

[1] Haider, M., Uhlemann, S.: Towards sub-ångstrom resolution with a 200 kv tem by means of a cscorrector and a computer controlled alignment procedure. Microscopy and Microanalysis 4(S2), 50–51 (1998)

[2] Yip, K.M., Fischer, N., Paknia, E., Chari, A., Stark, H.: Atomic-resolution protein structure determination by cryo-em. Nature 587(7832), 157–161 (2020)

[3] Nakane, T., Kotecha, A., Sente, A., McMullan, G., Masiulis, S., Brown, P.M., Grigoras, I.T., Malinauskaite, L., Malinauskas, T., Miehling, J., et al.: Single-particle cryo-em at atomic resolution. Nature 587(7832), 152–156 (2020)

[4] McMullan, G., Faruqi, A., Henderson, R.: Direct electron detectors. Methods in enzymology 579, 1–17 (2016)

[5] Maiden, A.M., Rodenburg, J.M.: An improved ptychographical phase retrieval algorithm for diffractive imaging. Ultramicroscopy 109(10), 1256–1262 (2009)

[6] Scheres, S.H.: Relion: implementation of a bayesian approach to cryo-em structure determination. Journal of structural biology 180(3), 519–530 (2012)

[7] Punjani, A., Rubinstein, J.L., Fleet, D.J., Brubaker, M.A.: cryosparc: algorithms for rapid unsupervised cryo-em structure determination. Nature methods 14(3), 290–296 (2017)

[8] Müller, K., Ryll, H., Ordavo, I., Ihle, S., Strüder, L., Volz, K., Zweck, J., Soltau, H., Rosenauer, A.: Scanning transmission electron microscopy strain measurement from millisecond frames of a direct electron charge coupled device. Applied Physics Letters 101(21), 212110 (2012) 10.1063/1.4767655

[9] Müller-Caspary, K., Oelsner, A., Potapov, P.: Two-dimensional strain mapping in semiconductors by nano-beam electron diffraction employing a delay-line detector. Applied Physics Letters 107(7) (2015)

[10] Clausen, A., Weber, D., Mueller, E., Poghosyan, E., Stroppa, D., Dunin-Burkowski, R.: Live data processing of 4D STEM experiments: LiberTEM meets ARINA hybrid-pixel detector. Oxford University Press US (2023)

[11] Rodenburg, J., Maiden, A.: Ptychography. Springer Handbook of Microscopy, 819–904 (2019)

[12] Nguyen, K.X., Jiang, Y., Lee, C.-H., Kharel, P., Zhang, Y., Zande, A.M., Huang, P.Y.: Achieving sub-0.5-angstrom–resolution ptychography in an uncorrected electron microscope. Science 383(6685), 865–870 (2024)

[13] Chen, Z., Jiang, Y., Shao, Y.-T., Holtz, M.E., Odstr čil, M., Guizar-Sicairos, M., Hanke, I., Ganschow, S., Schlom, D.G., Muller, D.A.: Electron ptychography achieves atomic-resolution limits set by lattice vibrations. Science 372(6544), 826–831 (2021)

[14] Li, G., Xu, M., Tang, W.-Q., Liu, Y., Chen, C., Zhang, D., Liu, L., Ning, S., Zhang, H., Gu, Z.-Y., et al.: Atomically resolved imaging of radiation-sensitive metal-organic frameworks via electron ptychography. Nature communications 16(1), 914 (2025)

[15] Yuan, B., Wang, Z., Zhang, S., Hofer, C., Gao, C., Chennit, T., Shi, H., Wu, X., Han, Y., Dou, L., et al.: Atomically resolved edges and defects in lead halide perovskites. Nature 647(8089), 364–368 (2025)

[16] Bennemann, F., Kirkland, A.I., Muller, D.A., Nellist, P.D.: Detective quantum efficiency-based comparison of hrtem and ptychography phase imaging. Microscopy and Microanalysis 32(2), 018 (2026)

[17] Zhou, L., Song, J., Kim, J.S., Pei, X., Huang, C., Boyce, M., Mendonca, L., Clare, D., Siebert, A., Allen, C.S., et al.: Low-dose phase retrieval of biological specimens using cryo-electron ptychography. Nature communications 11(1), 2773 (2020)

[18] Lazić, I., Wirix, M., Leidl, M.L., Haas, F., Mann, D., Beckers, M., Pechnikova, E.V., Müller-Caspary, K., Egoavil, R., Bosch, E.G., et al.: Single-particle cryo-em structures from idpc–stem at near-atomic resolution. Nature Methods 19(9), 1126–1136 (2022)

[19] Yu, Y., Spoth, K.A., Colletta, M., Nguyen, K.X., Zeltmann, S.E., Zhang, X.S., Paraan, M., Kopylov, M., Dubbeldam, C., Serwas, D., et al.: Dose-efficient cryo-electron microscopy for thick samples using tilt-corrected scanning transmission electron microscopy. Nature methods 22(10), 2138–2148 (2025)

[20] Seifer, S., Houben, L., Elbaum, M.: Shadow montage and cone-beam reconstruction in 4d-stem tomography. Microscopy and Microanalysis 32(1), 126 (2026)

[21] Rodenburg, J., McCallum, B., Nellist, P.: Experimental tests on double-resolution coherent imaging via stem. Ultramicroscopy 48(3), 304–314 (1993)

[22] Maiden, A.M., Humphry, M.J., Rodenburg, J.M.: Ptychographic transmission microscopy in three dimensions using a multi-slice approach. Journal of the Optical Society of America a 29(8), 1606–1614 (2012)

[23] Findlay, S.D., Brown, H.G., Pelz, P.M., Ophus, C., Ciston, J., Allen, L.J.: Scattering matrix determination in crystalline materials from 4d scanning transmission electron microscopy at a single defocus value. Microscopy and microanalysis 27(4), 744–757 (2021)

[24] Diederichs, B., Herdegen, Z., Strauch, A., Filbir, F., Müller-Caspary, K.: Exact inversion of partially coherent dynamical electron scattering for picometric structure retrieval. Nature Communications 15(1), 101 (2024)

[25] Küçükoğlu, B., Mohammed, I., Guerrero-Ferreira, R.C., Ribet, S.M., Varnavides, G., Leidl, M.L., Lau, K., Nazarov, S., Myasnikov, A., Kube, M., et al.: Low-dose cryo-electron ptychography of proteins at sub-nanometer resolution. Nature Communications 15(1), 8062 (2024)

[26] Pei, X., Zhou, L., Huang, C., Boyce, M., Kim, J.S., Liberti, E., Hu, Y., Sasaki, T., Nellist, P.D., Zhang, P., et al.: Cryogenic electron ptychographic single particle analysis with wide bandwidth information transfer. Nature Communications 14(1), 3027 (2023)

[27] Grant, T., Grigorieff, N.: Measuring the optimal exposure for single particle cryo-em using a 2.6 å reconstruction of rotavirus vp6. elife 4, 06980 (2015)

[28] Zheng, S.Q., Palovcak, E., Armache, J.-P., Verba, K.A., Cheng, Y., Agard, D.A.: Motioncor2: anisotropic correction of beam-induced motion for improved cryo-electron microscopy. Nature methods 14(4), 331–332 (2017)

[29] Li, X., Mooney, P., Zheng, S., Booth, C.R., Braunfeld, M.B., Gubbens, S., Agard, D.A., Cheng, Y.: Electron counting and beam-induced motion correction enable near-atomic-resolution single-particle cryo-em. Nature methods 10(6), 584–590 (2013)

[30] Zivanov, J., Nakane, T., Forsberg, B.O., Kimanius, D., Hagen, W.J., Lindahl, E., Scheres, S.H.: New tools for automated high-resolution cryo-em structure determination in relion-3. elife 7, 42166 (2018)

[31] Jamali, K., Käll, L., Zhang, R., Brown, A., Kimanius, D., Scheres, S.H.: Automated model building and protein identification in cryo-em maps. Nature 628(8007), 450–457 (2024)

[32] Varnavides, G., Ribet, S.M., Zeltmann, S.E., Yu, Y., Savitzky, B.H., Byrne, D.O., Allen, F.I., Dravid, V.P., Scott, M.C., Ophus, C.: Iterative phase retrieval algorithms for scanning transmission electron microscopy. arXiv preprint arXiv:2309.05250 (2023)

[33] Lupini, A.R.: The electron ronchigram. Scanning Transmission Electron Microscopy: Imaging and Analysis, 117–161 (2010)

[34] Lupini, A.R., Wang, P., Nellist, P.D., Kirkland, A.I., Pennycook, S.J.: Aberration measurement using the ronchigram contrast transfer function. Ultramicroscopy 110(7), 891–898 (2010) 10.1016/j.ultramic.2010.04.006

[35] Tegunov, D., Cramer, P.: Real-time cryo-electron microscopy data preprocessing with warp. Nature methods 16(11), 1146–1152 (2019)

[36] Elferich, J., Kong, L., Zottig, X., Grigorieff, N.: Ctffind5 provides improved insight into quality, tilt, and thickness of tem samples. Elife 13, 97227 (2024)

[37] Zhang, K.: Gctf: Real-time ctf determination and correction. Journal of Structural Biology 193(1), 1–12 (2016) 10.1016/j.jsb.2015.11.003

[38] Llopart, X., Alozy, J., Ballabriga, R., Campbell, M., Casanova, R., Gromov, V., Heijne, E., Poikela, T., Santin, E., Sriskaran, V., et al.: Timepix4, a large area pixel detector readout chip which can be tiled on 4 sides providing sub-200 ps timestamp binning. Journal of Instrumentation 17(01), 01044 (2022)

[39] Dushimineza, J.F., Leidl, M.L., Sturm, S., Denneulin, T., Müller-Caspary, K.: Imaging ionic displacements across inclined ferroelectric domains by a reciprocity approach to 4d-stem. Small in print, 75386 10.1002/smll.75386 https://onlinelibrary.wiley.com/doi/pdf/10.1002/smll.75386

[40] Jehle, A., Lorenzen, T., Leidl, M.L., Müller-Caspary, K.: The cramér-rao lower bound for the precision of scan position refinement in momentum-resolved stem. Ultramicroscopy 284, 114379 (2026) 10.1016/j.ultramic.2026.114379

[41] Imambi, S., Prakash, K.B., Kanagachidambaresan, G.: Pytorch. In: Programming with TensorFlow: Solution for Edge Computing Applications, pp. 87–104. Springer, ??? (2021)

[42] Rodenburg, J.M., Bates, R.H.T.: The Theory of Super-Resolution Electron Microscopy Via Wigner-Distribution Deconvolution. Phil. Trans. A: Phys. Sci. Engin. 339, 521–553 (1992) 10.1098/rsta.1992.0050

[43] Lalandec Robert, Hoelen L., Leidl, Max Leo, Müller-Caspary, Knut, Verbeeck, Jo: Benchmarking analytical electron ptychography methods for the low-dose imaging of beam-sensitive materials. Eur. Phys. J. Appl. Phys. 100, 20 (2025) 10.1051/epjap/2025018

[44] Leidl, M.L., Sachse, C., Müller-Caspary, K.: Dynamical scattering in ice-embedded proteins in conventional and scanning transmission electron microscopy. IUCrJ 10(4), 475–486 (2023)

[45] Leidl, M.L., Sturm, S., Filopoulou, A., Sachse, C., Müller-Caspary, K.: Towards a protein-size dependent resolution limit due to dynamical scattering in cryo-transmission electron microscopy. Microscopy and Microanalysis 31(6), 123 (2025)

[46] Akiba, T., Sano, S., Yanase, T., Ohta, T., Koyama, M.: Optuna: A next-generation hyperparameter optimization framework. In: Proceedings of the 25th ACM SIGKDD International Conference on Knowledge Discovery & Data Mining, pp. 2623–2631 (2019)

[47] Cao, M.C., Chen, Z., Jiang, Y., Han, Y.: Automatic parameter selection for electron ptychography via bayesian optimization. Scientific Reports 12(1), 12284 (2022)

[48] Noble, A.J., Dandey, V.P., Wei, H., Brasch, J., Chase, J., Acharya, P., Tan, Y.Z., Zhang, Z., Kim, L.Y., Scapin, G., et al.: Routine single particle cryoem sample and grid characterization by tomography. Elife 7, 34257 (2018)

[49] Fernandez-Gimenez, E., Carazo, J.M., Sorzano, C.: Local defocus estimation in single particle analysis in cryo-electron microscopy. Journal of structural biology 215(4), 108030 (2023)

[50] Bosch, E.G., Lazić, I.: Analysis of depth-sectioning stem for thick samples and 3d imaging. Ultramicroscopy 207, 112831 (2019)

[51] Lazić, I., Bosch, E.G., Lazar, S.: Phase contrast stem for thin samples: Integrated differential phase contrast. Ultramicroscopy 160, 265–280 (2016)

[52] Brilot, A.F., Chen, J.Z., Cheng, A., Pan, J., Harrison, S.C., Potter, C.S., Carragher, B., Henderson, R., Grigorieff, N.: Beam-induced motion of vitrified specimen on holey carbon film. Journal of structural biology 177(3), 630–637 (2012)

[53] Fromm, S.A., Bharat, T.A., Jakobi, A.J., Hagen, W.J., Sachse, C.: Seeing tobacco mosaic virus through direct electron detectors. Journal of structural biology 189(2), 87 (2015)

[54] Weis, F., Beckers, M., Der Hocht, I., Sachse, C.: Elucidation of the viral disassembly switch of tobacco mosaic virus. The EMBO Reports 20(11), 201948451 (2019)

[55] Meng, E.C., Pettersen, E.F., Couch, G.S., Huang, C.C., Ferrin, T.E.: Tools for integrated sequence-structure analysis with ucsf chimera. BMC bioinformatics 7(1), 339 (2006)

[56] Meng, E.C., Goddard, T.D., Pettersen, E.F., Couch, G.S., Pearson, Z.J., Morris, J.H., Ferrin, T.E.: Ucsf chimerax: Tools for structure building and analysis. Protein Science 32(11), 4792 (2023)

[57] Barad, B.A., Echols, N., Wang, R.Y.-R., Cheng, Y., DiMaio, F., Adams, P.D., Fraser, J.S.: Emringer: side chain–directed model and map validation for 3d cryoelectron microscopy. Nature methods 12(10), 943–946 (2015)

[58] Müller, K., Krause, F.F., Beche, A., Schowalter, M., Galioit, V., Löffler, S., Verbeeck, J., Zweck, J., Schattschneider, P., Rosenauer, A.: Atomic electric fields revealed by a quantum mechanical approach to electron picodiffraction. Nature Communications 5, 5653–18 (2014) 10.1038/ncomms6653

[59] Jannis, D., Hofer, C., Gao, C., Xie, X., Béché, A., Pennycook, T.J., Verbeeck, J.: Event driven 4d stem acquisition with a timepix3 detector: Microsecond dwell time and faster scans for high precision and low dose applications. Ultramicroscopy 233, 113423 (2022)

[60] Schayck, J.P., Zhang, Y., Knoops, K., Peters, P.J., Ravelli, R.B.: Integration of an event-driven timepix3 hybrid pixel detector into a cryo-em work-flow. Microscopy and Microanalysis 29(1), 352–363 (2023)

[61] Nakane, T., Kotecha, A., Sente, A., McMullan, G., Masiulis, S., Brown, P.M.G.E., Grigoras, I.T., Malinauskaite, L., Malinauskas, T., Miehling, J., Uchański, T., Yu, L., Karia, D., Pechnikova, E.V., Jong, E., Keizer, J., Bischoff, M., McCormack, J., Tiemeijer, P., Hardwick, S.W., Chirgadze, D.Y., Murshudov, G., Aricescu, A.R., Scheres, S.H.W.: Single-particle cryoem at atomic resolution. Nature 587(7832), 152–156 (2020) 10.1038/s41586-020-2829-0

[62] Haider, M., Uhlemann, S., Schwan, E., Rose, H., Kabius, B., Urban, K.: Electron microscopy image enhanced. Nature 392(6678), 768–769 (1998) 10.1038/33823

[63] Batson, P.E., Dellby, N., Krivanek, O.L.: Sub-angstrom resolution using aberration corrected electron optics. Nature 418, 617–620 (2002) 10.1038/nature00972

[64] Lorenzen, T., März, B., Xue, T., Beyer, A., Volz, K., Bein, T., Müller-Caspary, K.: Imaging built-in electric fields and light matter by fourier-precession tem. Scientific Reports 14, 1320 (2024) 10.1038/s41598-024-51423-x

[65] Yuan, B., Wang, Z., Zhang, S., Hofer, C., Gao, C., Chennit, T., Shi, H., Wu, X., Han, Y., Dou, L., Yu, Y., Pennycook, T.J.: Atomically resolved edges and defects in lead halide perovskites. Nature 647(8089), 364–368 (2025) 10.1038/s41586-025-09693-6

[66] Lučić, V., Rigort, A., Baumeister, W.: Cryo-electron tomography: the challenge of doing structural biology in situ. Journal of Cell Biology 202(3), 407–419 (2013)

[67] Romanov, A., Cho, M.G., Scott, M.C., Pelz, P.: Multi-slice electron ptychographic tomography for three-dimensional phase-contrast microscopy beyond the depth of focus limits. JPhys Materials 8 (2025) 10.1088/2515-7639/ad9ad2 . CRIS-Team Scopus Importer:2025-03-14

[68] Pelz, P., Griffin, S.M., Stonemeyer, S., Popple, D., DeVyldere, H., Ercius, P., Zettl, A., Scott, M.C., Ophus, C.: Solving complex nanostructures with ptychographic atomic electron tomography. Nature Communications 14 (2023) 10.1038/s41467-023-43634-z. CRIS-Team Scopus Importer:2023-12-15

[69] You, S., Romanov, A., Pelz, P.: Near-isotropic sub-Ångstrom 3d resolution phase contrast imaging achieved by end-to-end ptychographic electron tomography. Physica Scripta 100, 015404 (2024) 10.1088/1402-4896/ad9a1a

[70] Thörnig, P.: Jureca: Data centric and booster modules implementing the modular supercomputing architecture at jülich supercomputing centre. Journal of large-scale research facilities JLSRF 7, 182–182 (2021)

[71] Lorenzen, T., Diederichs, B., Otieno Ogolla, C., Butz, B., Müller-Caspary, K.: Aberration measurement by electron ptychography and consistency among different algorithms. Applied Physics Letters 126(8) (2025)

[72] Müller-Caspary, K., Oppermann, O., Grieb, T., Krause, F.F., Rosenauer, A., Schowalter, M., Mehrtens, T., Beyer, A., Volz, K., Potapov, P.: Materials characterisation by angle-resolved scanning transmission electron microscopy. Scientific reports 6(1), 37146 (2016)

[73] YAML Ain’t Markup Language. https://yaml.org. Accessed: 2026-08-27

[74] Haider, M., Uhlemann, S., Zach, J.: Upper limits for the residual aberrations of a high-resolution aberration-corrected stem. Ultramicroscopy 81(3-4), 163–175 (2000)

[75] Rohou, A., Grigorieff, N.: Ctffind4: Fast and accurate defocus estimation from electron micrographs. Journal of structural biology 192(2), 216–221 (2015)

[76] Liu, D.C., Nocedal, J.: On the limited memory bfgs method for large scale optimization. Mathematical programming 45(1), 503–528 (1989)

[77] Agarap, A.F.: Deep learning using rectified linear units (relu). arXiv preprint arXiv:1803.08375 (2018)

[78] Guizar-Sicairos, M., Thurman, S.T., Fienup, J.R.: Efficient image registration algorithms for computation of invariant error metrics. In: Signal Recovery and Synthesis, p. 3 (2007). Optica Publishing Group

[79] Watanabe, S.: Tree-structured parzen estimator: Understanding its algorithm components and their roles for better empirical performance. arXiv preprint arXiv:2304.11127 (2023)

[80] Li, L., Jamieson, K., DeSalvo, G., Rostamizadeh, A., Talwalkar, A.: Hyperband: A novel bandit-based approach to hyperparameter optimization. Journal of machine learning research 18(185), 1–52 (2018)

[81] Cheng, A., Henderson, R., Mastronarde, D., Ludtke, S.J., Schoenmakers, R.H., Short, J., Marabini, R., Dallakyan, S., Agard, D., Winn, M.: Mrc2014: Extensions to the mrc format header for electron cryomicroscopy and tomography. Journal of structural biology 192(2), 146–150 (2015)

[82] Zivanov, J., Nakane, T., Scheres, S.H.: A bayesian approach to beam-induced motion correction in cryoem single-particle analysis. IUCrJ 6(1), 5–17 (2019)

[83] Böttcher, B., Nassal, M.: Structure of mutant hepatitis b core protein capsids with premature secretion phenotype. Journal of molecular biology 430(24), 4941–4954 (2018)

