## Supplementary Information for "Robust parameter estimation and dose-fractionation for ptychographic single-particle analysis at near-atomic resolution"

---

#### Supplementary Note 1: Compensating scan-induced beam tilt

In a STEM, the scan is realised by sets of coils that introduce a tilt and subsequent de-tilt of the beam, respectively. Especially when scanning large fields of view as in single-particle analysis, residual smooth variations of the incident beam direction across the scan remain. This translates directly to a scan position dependent shift of the bright field disc as a whole in diffraction space. Since ptychographic reconstructions rely on minimising the discrepancy between experimental and model diffraction patterns via their pixel-wise comparison, this effect is critical for the accurate retrieval of both the probe wave function and in particular the projected potential of the specimen.

We provide two methods to measure the bright field position in dependence of the scan position. Firstly, when the movement can be assumed to be constant throughout a session, we average the centre of mass (COM) of all diffraction patterns acquired at the same pixel of the scan grid and perform a polynomial fit to the COM. Secondly, when the acquisition parameters changed between subsequent scans, we propose a technique based on minimising the mismatch between the diffraction patterns at each point and their average. The developed algorithm is shown schematically in Suppl. Fig. 1 a and consists of two nested loops. In the outer loop depicted on the left an aligned position-averaged CBED (PACBED) is calculated using the currently estimated bright field positions, and its cross-correlations with all diffraction patterns are calculated. In the inner loop (dashed box) a set of polynomial coefficients is optimised to maximise the sum of cross correlations sampled at the predicted bright field positions.

The resulting bright field shifts relative to the optical axis are exemplarily shown for a scan across a vitrified apoferritin sample in Suppl. Fig. 1 b. The scan-induced movement of the bright field disc is significant and amounts to $\pm 2.2$ and $\pm 1.5$ pixels or equivalently $\pm 0.29$ and $\pm 0.20\,\mathrm{mrad}$ in both directions. The impact of the scan-induced beam tilt correction is demonstrated in Suppl. Fig. 1 c-e. Whereas the raw PACBED on the left in Suppl. Fig. 1 c shows significant smearing, especially in the vertical direction, a much clearer PACBED is obtained after aligning as displayed on the right. The Fresnel fringes appearing in the aligned pattern are a result of the Fringe-Free-Imaging (FFI) alignment of the microscope dedicated for plane wave illumination TEM [2], causing Fresnel fringes by an electron-optically displaced probe-forming aperture which we incorporated as a near-field Fresnel diffraction effect.

With defocus, astigmatism and scanning geometry initialised as described above for the scan across apoferritin, probe and specimen have been retrieved ptychographically as shown in Suppl. Fig. 1 d and e, respectively. The left parts depict a reconstruction in which the movement of the bright field disc was ignored, whereas the measured positions from Suppl. Fig. 1 b were incorporated for the right hand side. During the reconstruction, we treat the tilt in first order by applying a phase wedge to the scanning probe in real space before interaction with the specimen, assuming that the effect of aberrations is invariant with respect to the scan position. In fact, this improves the reconstructions in that the probe on the right in Suppl. Fig. 1 d shows the concentric rings expected for a defocused probe while the probe contains severe artefacts. This increase in probe quality also translates into enhanced phase reconstructions. Whereas apoferritin particles are difficult to detect in the standard reconstruction on the left in Suppl. Fig. 1 e, our procedure successfully recovers the apoferritin particles at high contrast in the right half. The particles are visible across the full field of view as highlighted by the zoomed-in patches shown in Suppl. Fig. 1 f.

Experimentally, both methods yield comparable results for the non-dose-fractionated experiments. However, the second, iterative approach can fail in ultimate low-dose situations, e.g., when the SNR of a single scan suffers from fractionating the total dose across multiple 4D-STEM recordings. While the low-dose sensitivity can be partially reduced by binning the dataset in the scan dimension if the movement between neighbouring scan positions is rather small, the COM-based, first method was employed in this work because of its robustness and low computational cost.

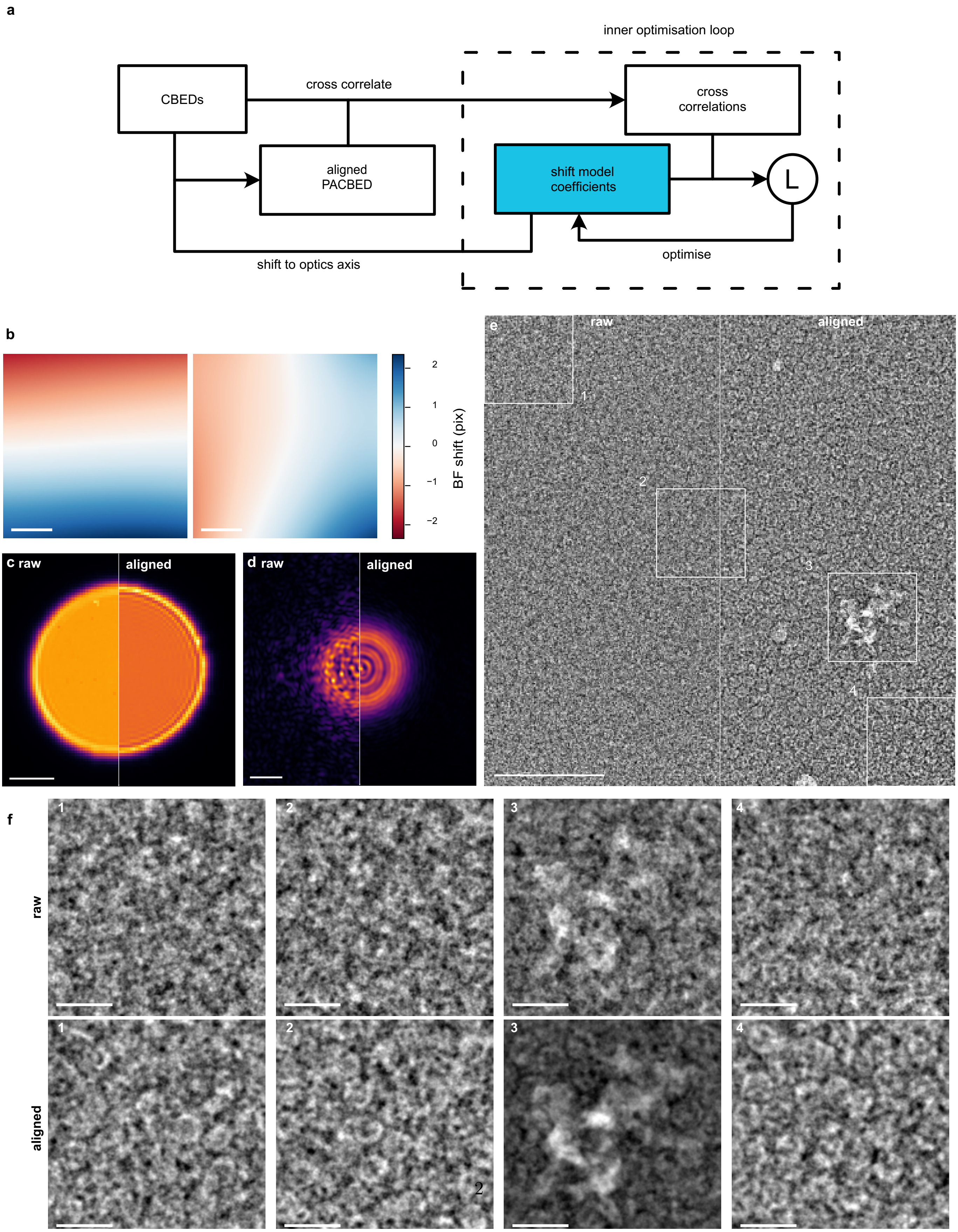


**Supplementary Figure 1 Estimation of bright field movement and its effect on reconstructions.** (a) Algorithm for the estimation of bright field movement. In the inner loop the shift is optimised by maximising the score L, in the outer loop an aligned PACBED is calculated. (b) Determined shifts in $q_x$- and $q_y$-direction (100 nm scalebar). (c) Raw and aligned mean diffraction pattern (2 mrad scalebar). (d) Electron probe (2 nm scalebar) and (e) object phase (100 nm scalebar) reconstructed with and without eliminating the shifting bright field disk in the reconstruction. (f) The highlighted specimen regions in (e) are shown lowpass-filtered in (20 nm scalebar). The full images are shown in Suppl. Fig. 12.

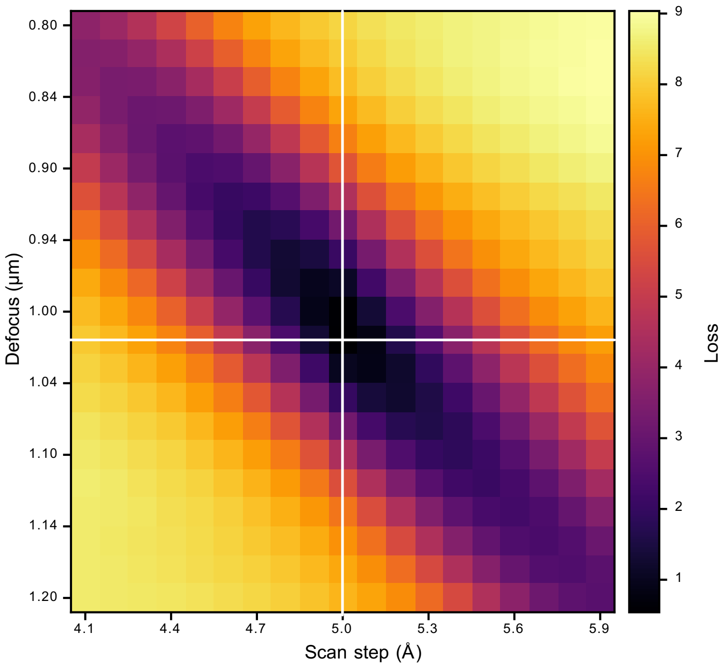


**Supplementary Figure 2 Entanglement of scan step and defocus.** Reconstruction loss of simulated 4D-STEM data plotted against the used defocus and scan step. When reconstructing with a scan step that deviates from the ground truth (white line, 5 Å) a wrong defocus is likely obtained.

**Supplementary Table 1 Acquisition parameters.** Parameters for datasets appearing in the manuscript.

| | Fig. 2 c-e, 3, Suppl. Fig. 4 a, 3 | Fig. 2 f-h, Suppl. Fig. 4 b-e | Suppl. Fig. 1 |
|---|---|---|---|
| Microscope | Titan Krios EPFL | Titan Themis 80-300 | Titan Krios ER-C |
| High tension (kV) | 300 | 200 | 300 |
| Semi-convergence angle (mrad) | 6 | 8.9 | 4 |
| Spherical aberration (mm) | 0. | 0. | 2.7 |
| Detector | Dectris Arina | QD MerlinT4 | Dectris Arina CZT |
| Scan size | 250 x 250 | 256 x 256 | 1024 x 1024 |
| Scan pixel size (Å) | 20 | 20 | 4.96 |
| Dose ($e^-$ / Å$^2$) | 35 | 20 | 25 |

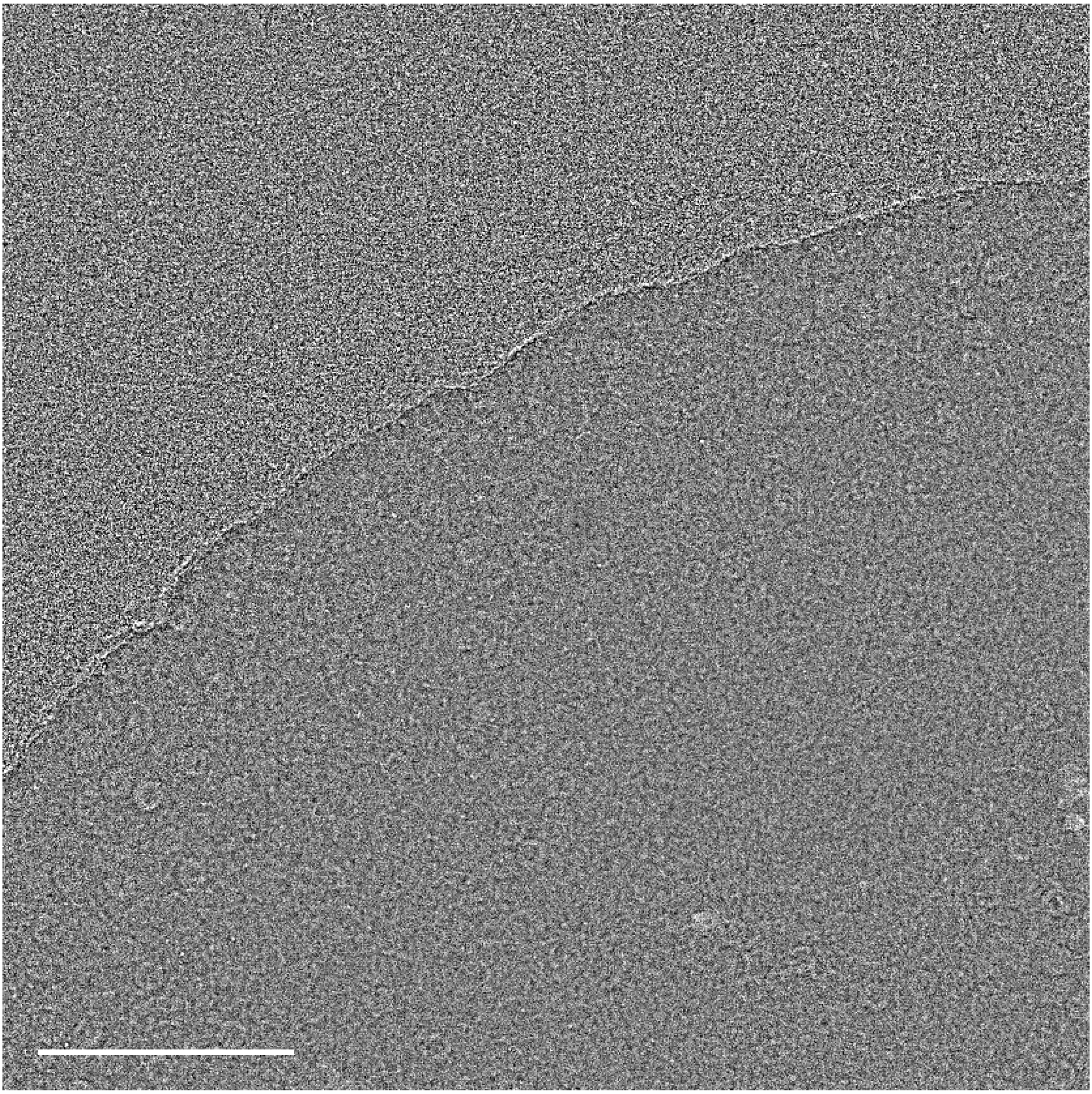

**Supplementary Figure 3 Apoferritin micrograph.** Ptychographic reconstruction performed using the non-tilted dataset presented in Fig. 3, the length of the scalebar equals 100 nm.

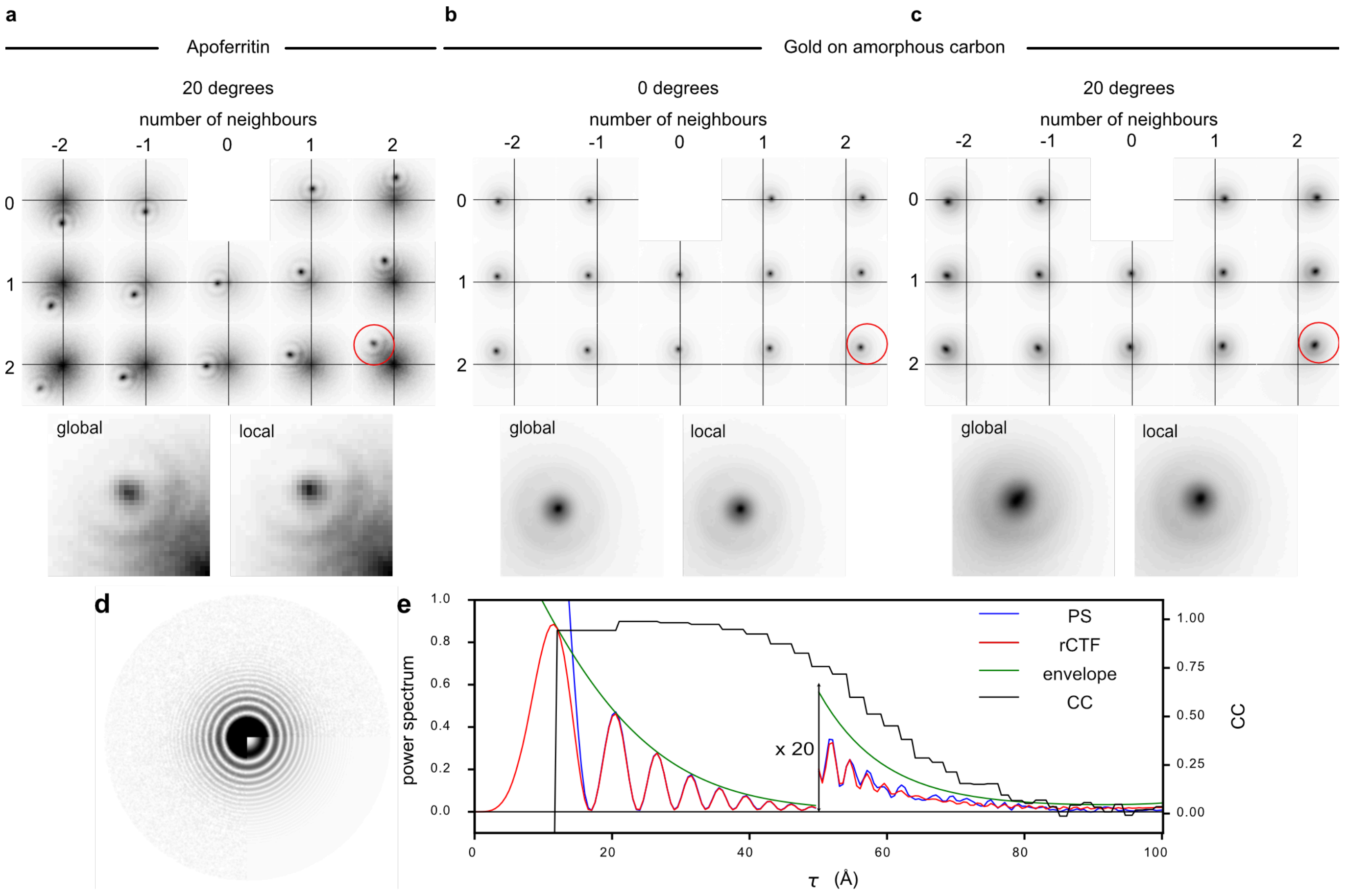


**Supplementary Figure 4 Scan parameter estimation using the Ronchigram cross-correlation.** Ronchigram cross-correlation plots for (a) a 20 degrees tilted apoferritin grid, (b) a non-tilted and (c) a 20 degree tilted gold on amorphous carbon specimen. The numbers on the top and side indicate neighbour orders in the fast and slow scan direction, respectively. Shown are the central parts of the neighbour correlation up to two neighbours away in both directions. The black crosses indicate the center of the coordinate system. The bottom shows the peak of a cross-correlation (highlighted by the red circle) before (global) and after (local) rescaling using the local defocus distribution from Fig. 2. Only a small increase in peak quality is observed for the non-tilted specimen while peak quality strongly increases in the tilted case. (d) Power spectrum (square root scale) of the shadow image recovered from the cross-correlation peaks in (b) and the fitted rCTF. (e) Azimuthal average of the same area.

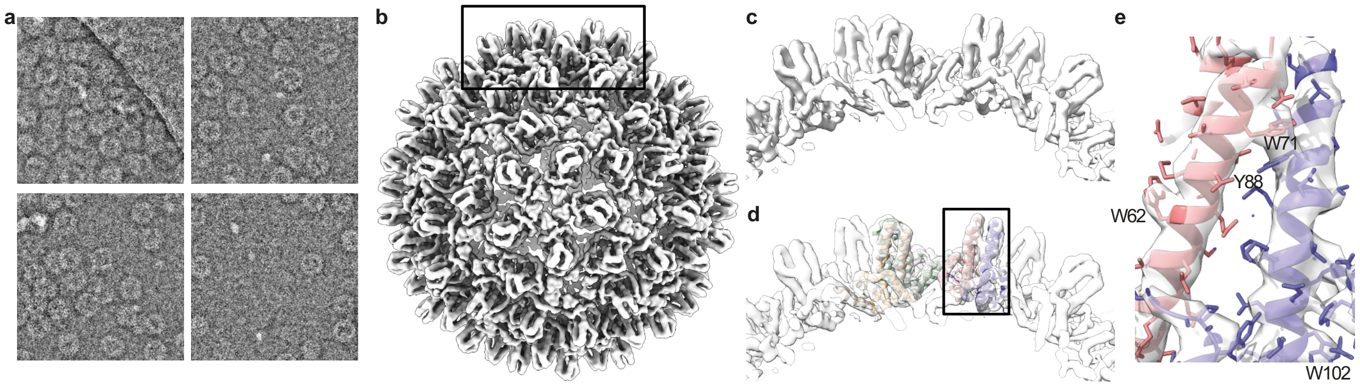


**Supplementary Figure 5  single-particle 3D-reconstruction of DNA-filled HBV capsids at 4.5 Å resolution.** (a) Ptychographic micrographs of DNA-filled HBV capsids. (b) Full capsid and zoomed spikes (black box in b) (c) without and (d) with docked atomic model (fitted model ID: PDB 6HTX) [83]. (e) Zoomed spike (black box in d) showing bulky side chains.

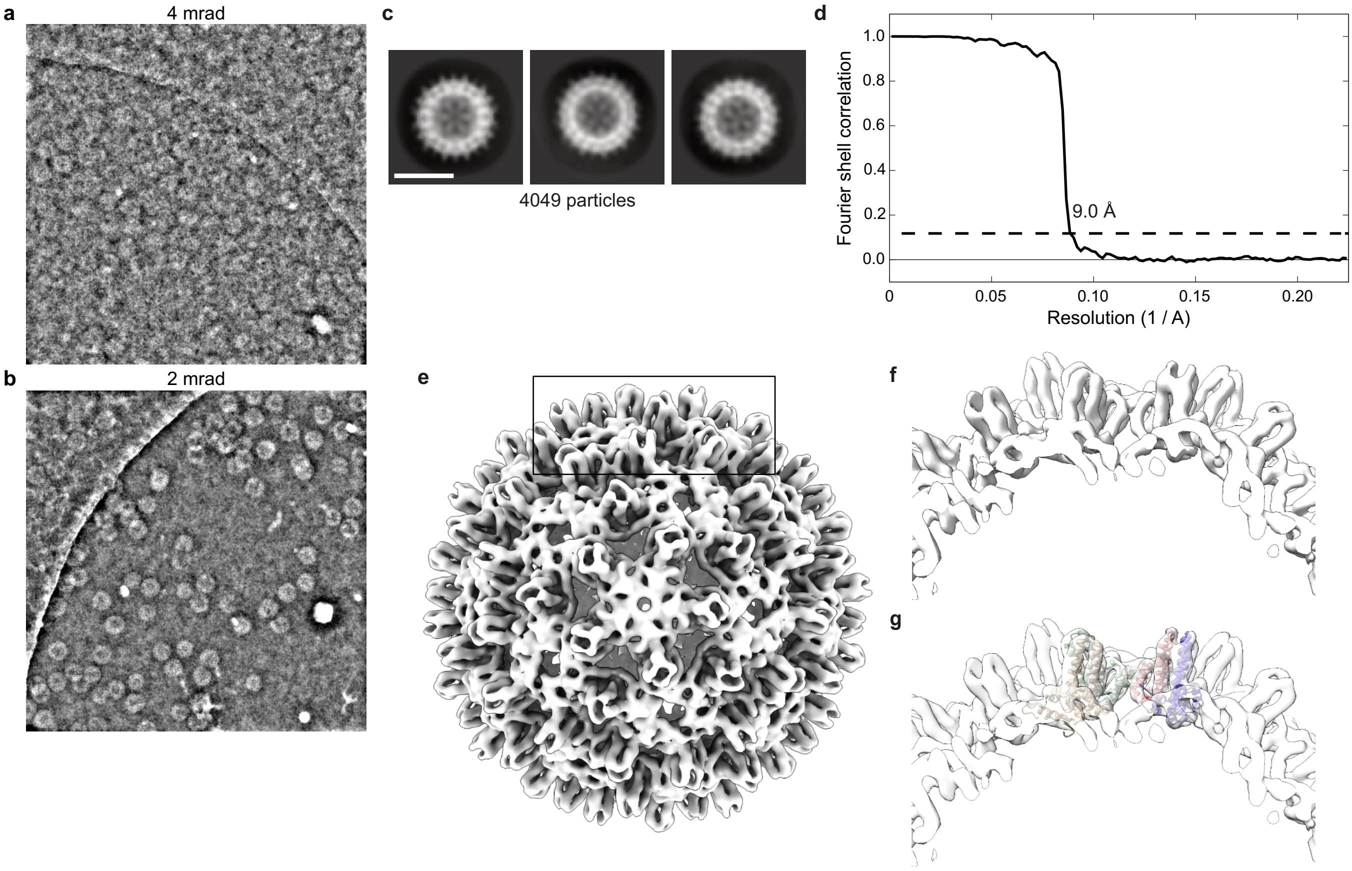


**Supplementary Figure 6  iDPC-STEM imaging of HBV capsids.** iDPC micrographs of (a) HBV capsids (RNA-filled) at 4 mrad SCA and (b) HBV capsids (DNA-filled) at 2 mrad SCA with strongly increased contrast in (b). (c) 2D-classes selected for further processing. (d) FSC curve of the 3D-reconstruction demonstrating 9.0 Å resolution. (e-g) Reproduction of Suppl. Fig. 5 using the iDPC map. The length of the scalebar in (c) equals 23 nm.

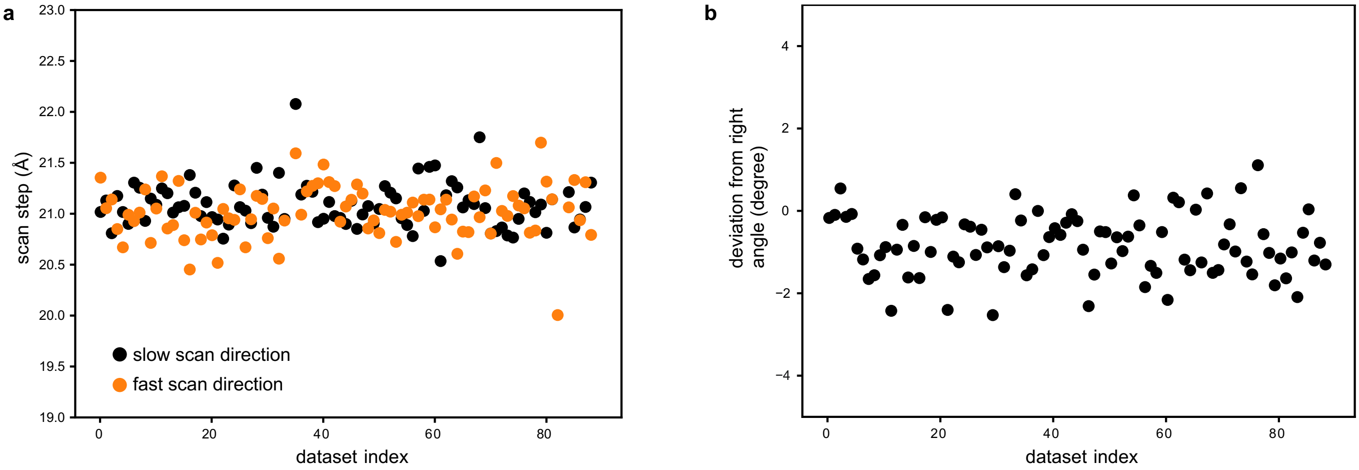


**Supplementary Figure 7 Scanning geometry estimation in EMPIAR-12236.** (a) Length of the scan step in the fast and slow scanning direction, respectively, as determined using the Ronchigram cross-correlation. (b) Deviation of the angle between the slow and fast scanning step from 90 degrees.

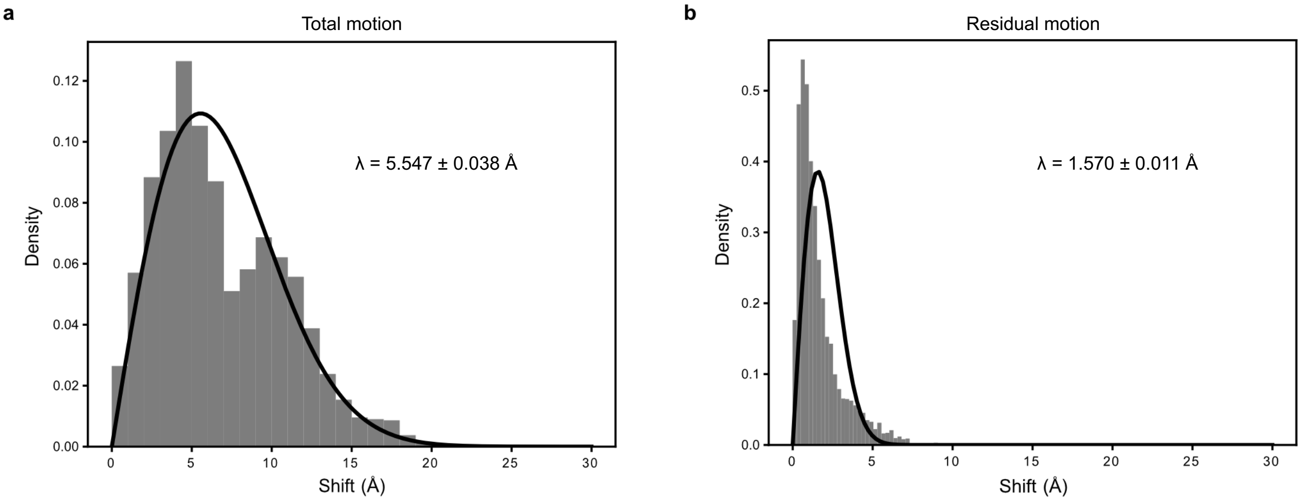


**Supplementary Figure 8 Particle motion.** Histogram of particle shifts observed in the dose-fractionated TMV datasets (a) including the full-frame motion and (b) after removal of the full-frame motion. The mean shifts $\lambda$ and plotted Rayleigh distributions (black lines) were determined according to Ref. [1]. The distribution in (b) strongly deviates from the fitted Rayleigh distribution. One possible source is the subtracted rigid motion which not only includes the stage movement but also the joint movement of the particles. This joint movement is a large part of the particle motion due to the small FOV in the reconstructions of ca. 250 x 250 $\text{nm}^2$. Additionally, the motion of neighbouring particles is highly correlated along the rods.

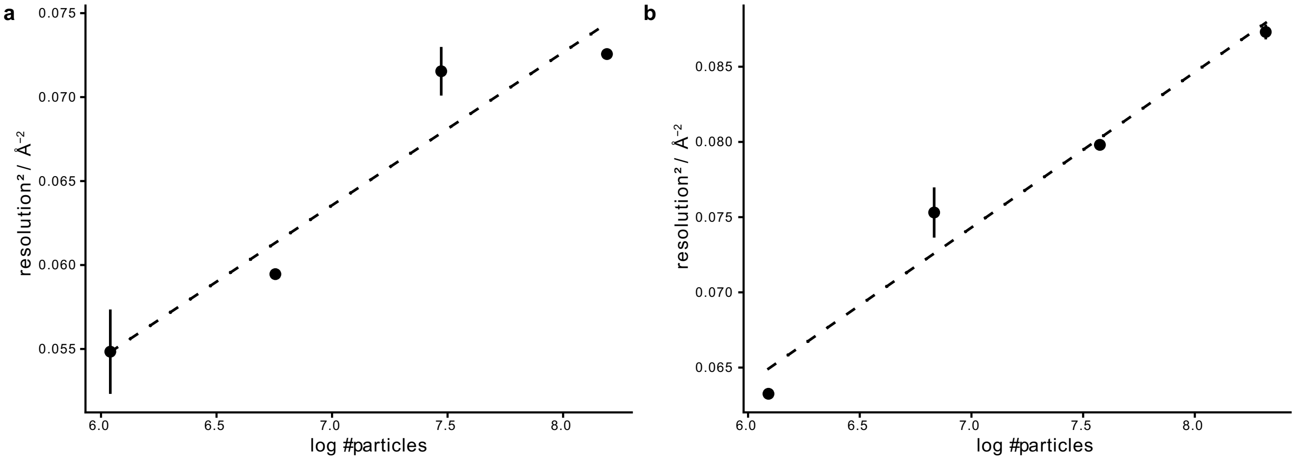


**Supplementary Figure 9 Calculation of ResLog B-Factors.** ResLog plot for (a) standard ptychographic SPA for TMV and (b) dose-fractionated SPA on TMV. Helical 3D-reconstructions were performed for three random subsets of particles at each particle number.

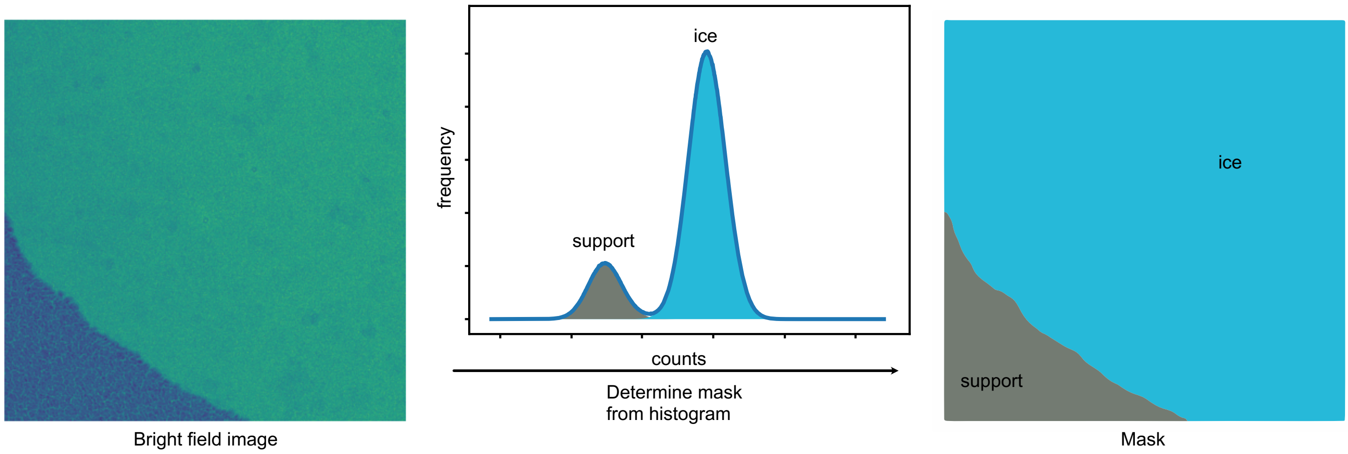


**Supplementary Figure 10 Masking of support.** Utilising histograms of bright field images (left), scan regions belonging to the strongly scattering support are identified and removed from further processing.

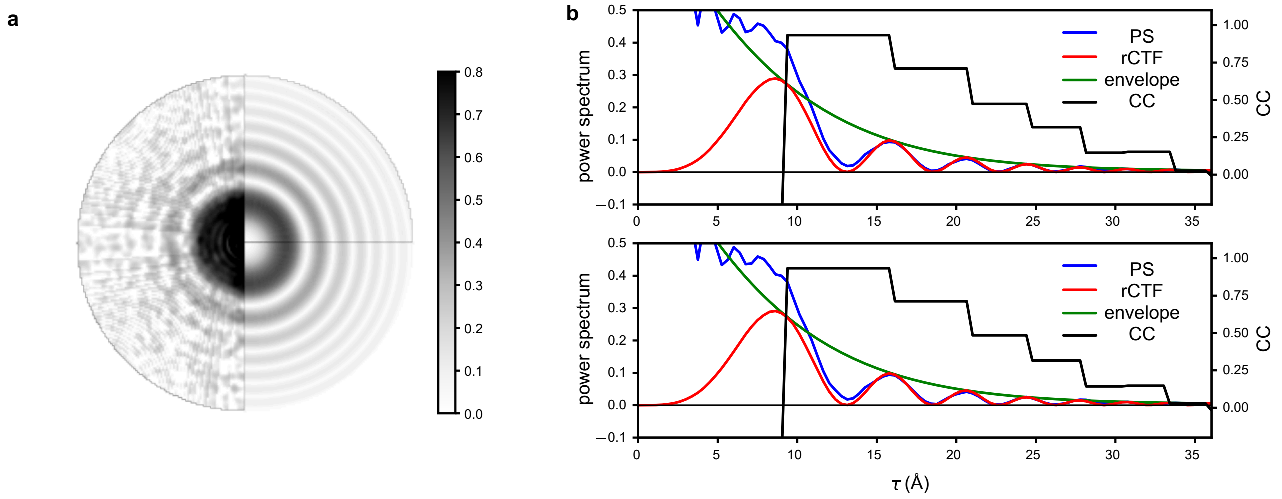


**Supplementary Figure 11 Influence of the envelope on the rCTF determination.** During rCTF determination defocus and astigmatism are first estimated by maximising the normalised correlation of the background-subtracted square root of the power spectrum ("amplitude spectrum") with the rCTF model, neglecting any envelope effects. This result is refined in a second fitting step on the full power spectrum using a parameterised background and envelope model. While the inclusion of the envelope significantly increases agreement with the model, the effects on the fitted rCTF parameters are minimal. Using the initial estimation without an envelope a defocus of 873 nm and a complex astigmatism of 6+53i nm was determined, while the refinement including the envelope resulted in minimal change, producing a defocus of 876 nm and a complex astigmatism of 7+56i nm. (a) The background-subtracted power spectrum (square root scale), together with an rCTF model including the effect of the envelope (top right) and the initially estimated rCTF model (bottom right). (b) The azimuthal averages using the refined (top) and initial rCTF model (bottom).

**Supplementary Table 2 Statistics of the presented ptychographic SPA datasets.**

| | TMV | TMV, fractionated | Apoferritin | HBV capsids | Mouse ApoF (EMPIAR-12236) |
|---|---|---|---|---|---|
| Microscope | TFS Titan Krios G4 ER-C, Dectris Arina CZT | | | | TFS Titan Krios G4<br>Cs probe corrector<br>Dectris ELA |
| High tension (kV) | 300 | 300 | 300 | 300 | 300 |
| Step size (Å) | 4.96 | 4.96 | 4.96 | 5.04 | 20 |
| Semi-convergence angle (mrad) | 4 | 4 | 4 | 4 | 4 |
| Probe positions | 1024 x 1024 | 3 x 1024 x 1024 | 1024 x 1024 | 1024 x 1024 | 128 x 128 |
| Electron dose ($e^-$ / $Å^2$) | 25 | 23 | 25 | 25 | 35 |
| Specimen positions | 30 | 15 | 80 | 108 | 90 |
| Defocus (nm) | −190 – −800<br>270 – 1200 | −260 – −530 | −220 – −920<br>200 – 660 | −250 – −840<br>380 – 940 | −2550 – −1550 |
| Pixel size (Å) | 0.816 | 0.871 | 0.794 | 0.775 | 0.730 |
| Particle numbers | 3,607 | 5,285 | 4,999 | 4,049 | 22,198 |
| Number of asymmetric units | 504,980 | 739,900 | 119,976 | 242,940 | 532,752 |
| Symmetry | helical | | O | I | O |
| Map resolution (Å) | 3.7 | 3.3 | 4.1 | 4.5 | 3.7 |
| Guinier B-factor ($Å^2$) | 100 | 76 | 190 | 304 | 165 |
| ResLog B-factor ($Å^2$) | 220 | 194 | - | - | - |

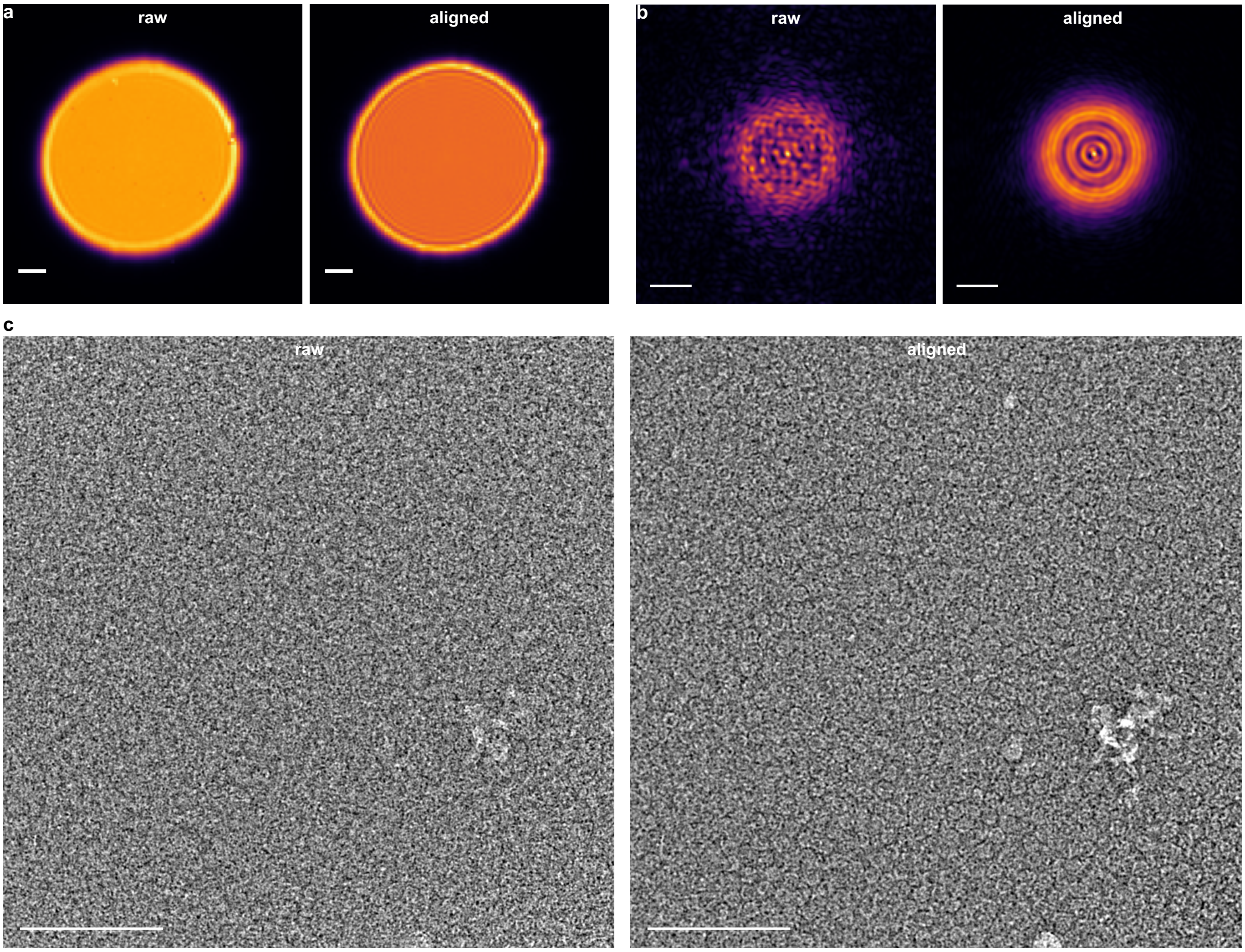


**Supplementary Figure 12 Estimation of bright field movement.** Mean diffraction pattern (a) without (raw) and (b) with compensation of the bright field movement (aligned, length of scalebar: 1 mrad). (b) Electron probe and (c) ptychographic micrograph obtained from a reconstruction without (raw) and with elimination of the bright field disk shift (aligned). The length of the scalebar in (b) is 2 nm and in (c) 100 nm.

### Supplementary References

[1] Axel F Brilot, James Z Chen, Anchi Cheng, Junhua Pan, Stephen C Harrison, Clinton S Potter, Bridget Carragher, Richard Henderson, and Nikolaus Grigorieff. Beam-induced motion of vitrified specimen on holey carbon film. *Journal of structural biology*, 177(3):630–637, 2012.

[2] Felix Weis and Wim JH Hagen. Combining high throughput and high quality for cryo-electron microscopy data collection. *Biological Crystallography*, 76(8):724–728, 2020.
